# Within population plastic responses to combined thermal-nutritional stress differ from those in response to single stressors, and are genetically independent across traits in both males and females

**DOI:** 10.1101/2023.03.26.534290

**Authors:** Yeuk Man Movis Choy, Greg M. Walter, Christen K. Mirth, Carla M. Sgrò

## Abstract

Phenotypic plasticity helps animals to buffer the effects of increasing thermal and nutritional stress created by climate change. Plastic responses to single and combined stressors can vary among genetically diverged populations. However, less is known about how plasticity in response to combined stress varies among individuals within a population or whether such variation changes across life-history traits. This is important because individual variation within populations shapes population-level responses to environmental change. Here, we used isogenic lines of *Drosophila melanogaster* to assess plasticity of egg-to-adult viability and sex-specific body size for combinations of two temperatures (25°C or 28°C) and three diets (standard diet, low caloric diet, or low protein:carbohydrate ratio diet). Our results reveal substantial within-population genetic variation in plasticity for egg-to-adult viability and wing size in response to combined thermal-nutritional stress. This genetic variation in plasticity was a result of cross-environment genetic correlations that were often < 1 for both traits, as well as changes in the expression of genetic variation across environments for egg-to-adult viability. Cross-sex genetic correlations for body size were weaker when the sexes were reared in different conditions, suggesting that the genetic basis of traits may change with the environment. Further, our results suggest that plasticity in egg-to-adult viability is genetically independent from plasticity in body size. Importantly, plasticity in response to diet and temperature individually differed from plastic shifts in response to diet and temperature in combination. By quantifying plasticity and the expression of genetic variance in response to combined stress across traits, our study reveals the complexity of animal responses to environmental change, and the need for a more nuanced understanding of the potential for populations to adapt to ongoing climate change.

## Introduction

Rises in temperatures and increased frequency of extreme weather events caused by global change are placing many organisms at a higher risk of extinction (Diffenbaugh & Field, 2013; Midgley & Hannah, 2019; Wiens, 2016). In addition to exposing organisms to more stressful thermal environments, climate change is also creating shifts in precipitation, which in combination with the increasing CO_2_ concentration, are affecting the distribution, abundance (Pecl et al., 2017; Van Der Putten et al., 2010), and nutritional composition of plants (Jin et al., 2019; Kreuzwieser & Gessler, 2010; Lukac et al., 2010). Specifically, plants are expected to contain less protein and varied carbohydrate content under climate change (Bowes, 1993; DaMatta et al., 2010; Dwyer et al., 2007; Richardson et al., 2002; Rosenblatt & Schmitz, 2016), potentially imposing direct nutritional stress on herbivores and frugivores (Bowes, 1993; DaMatta et al., 2010; Dwyer et al., 2007; Richardson et al., 2002; Rosenblatt & Schmitz, 2016) and indirectly through changes in yeast communities. In addition, ectothermic herbivores and frugivores will be at a greater risk as they rely on ambient temperatures to maintain their metabolism (Hawlena & Schmitz, 2010; Walters et al., 2012). As one of the largest groups of primary consumers, small changes in the population dynamics of these ectotherms could affect the whole food chain and disturb ecosystem function (Rosenblatt & Schmitz, 2016). Understanding how ectothermic frugivores respond to combined thermal and nutritional stress will be crucial in predicting how species and ecosystems will respond to ongoing climate change.

Phenotypic plasticity – the ability of a single genotype to express different phenotypes in different environments – is often the first response of many organisms to environmental variation (Huey et al., 2002). Plastic shifts in life-history traits in response to environmental change are common (Abram et al., 2017; Schilthuizen & Kellermann, 2014; Seebacher et al., 2015; Sgrò et al., 2016; Urban et al., 2014; Via et al., 1995). While plasticity might contribute to persistence and adaptation to environmental change, it may not always be adaptive (Ghalambor et al., 2007). In small ectotherms such as insects, many studies have revealed plastic shifts in response to either temperature or nutrition. For instance, body size (Bochdanovits & De Jong, 2003; French et al., 1998), development time (Schou et al., 2017), and reproductive success (Janowitz & Fischer, 2011; S. Wang et al., 2013) have been shown to decrease with increasing temperature. Pre-adult viability (Jang & Lee, 2018; Min et al., 2007), development time (Economos & Lints, 1984), body size (Jang & Lee, 2018; Min et al., 2007), and reproductive success (Bader & Williams, 2012) also shift plastically in response to reductions in food quality (protein and carbohydrate contents) and quantity. However wild animals are exposed to multiple stressors simultaneously (Rosenblatt & Schmitz, 2016; Stillwell et al., 2007), which makes it important to understand plastic shifts in response to combinations of environmental stressors predicted to occur under climate change.

Recent studies focussed on understanding plastic responses to combinations of stressors have shown that trait responses to combined stress can change in unpredictable ways when compared to single stressor responses (Bubliy et al., 2012; Chakraborty et al., 2020; Kutz et al., 2019; Polak et al., 2004). Plastic responses to combined thermal-nutritional stress in insects have been explored for a range of life-history traits, including viability, developmental time, body size, and lifespan (Clissold & Simpson, 2015; Couret et al., 2014; Kutz et al., 2019; Rho & Lee, 2017; Tun-Lin et al., 2000). For example, in adult mealworm beetles (*Tenebrio molitor*), higher temperatures significantly reduced survival on sub-optimal diets, whereas diet did not impact survival at a lower temperature (Rho & Lee, 2017). Similarly, in *D. melanogaster*, the negative impacts of sub-optimal diets on viability were exacerbated at higher temperatures (Kutz et al., 2019). These studies suggest that changes in temperature and nutrition may interact to pose greater levels of stress than they do individually.

Plastic responses to environmental change can vary among populations and between individuals within populations, quantified as genotype by environment interactions (G × E) (Chakraborty et al., 2023; Dreyer et al., 2016; Frankino et al., 2019; Via & Lande, 1985). Several studies have looked at among-population variation in plasticity along latitudinal gradients (Chakraborty et al., 2020; Schmidt et al., 2005; Sgrò et al., 2010; Sisodia & Singh, 2010), while fewer studies have quantified genetic variation in plasticity within a population or between individual genotypes to thermal and nutritional stressors (Chakraborty et al., 2023; Frankino et al., 2019; Lafuente et al., 2018; Mossman et al., 2016; J. B. Wang et al., 2017). Variation in plasticity within a population provides the potential for plasticity to evolve at a population level, which will determine population responses to environmental change (Frankino et al., 2019; Ghalambor et al., 2007; Lande, 2009). Greater within-population genetic variation in plasticity will therefore increase the potential for plasticity to evolve, which will allow populations to adapt to environmental change (Chevin & Hoffmann, 2017; Lande, 2009; Walter et al., 2023). Understanding the distribution of plastic responses (shape of reaction norms) between individuals of a population is therefore vital for determining whether populations will persist under climate change.

A powerful approach for measuring within-population variation in plasticity is to create isogenic lines, which involves inbreeding lines that are collected from a natural population with each resulting isogenic line representing a fixed genotype (Franěk et al., 2020; Ørsted et al., 2018). Variation among isogenic lines within environments reflects snapshots of standing genetic variation of the sampled population. By measuring differences in the response of isogenic lines to multiple environments, we can quantify variation in plasticity among genotypes (Scheiner, 1993; Via & Conner, 1995). Variation in plasticity among genotypes (or significant G × E) can be caused by differences in the expression of genetic variation across environments, or cross-environment genetic correlations that are less than one (Via & Conner, 1995), or both. Understanding how the expression of genetic variation and genetic correlations change across environment and contribute to within-population variation in plasticity is essential for predicting a population’s ability to adapt to long-term environmental change (Ghalambor et al., 2007; Sgrò & Hoffmann, 2004; Wood & Brodie, 2015).

In this study, we used a set of newly derived isogenic lines of *D. melanogaster* initiated from a population collected from the east coast of Australia (Chakraborty et al., 2023). These lines allow us to extend upon previous studies that have examined variation in plasticity between populations sampled from the latitudinal cline along the east Australian coast (Chakraborty et al., 2020; Cockerell et al., 2014; Sgrò et al., 2010; van Heerwaarden et al., 2016). Specifically, we used these lines to quantify sex-specific variation in plasticity in response to combined thermal and nutritional stress. We subjected developing larvae to six combinations of thermal and nutritional conditions that included two temperatures: 25°C and 28°C, and three diets: standard, low caloric, and low protein to carbohydrate (P:C) diets. These conditions were chosen to best simulate climate change, which is expected to cause a 3°C increase in average temperature (Screen, 2014), and to reduce protein and vary carbohydrate contents in plants (DaMatta *et al*. 2010; Gowik and Westhoff 2011; IPCC 2018). We assessed plasticity of egg-to-adult viability and adult wing centroid size, which have previously shown sensitivity to combined thermal and nutritional stress (Chakraborty et al., 2020; Kutz et al., 2019). Given that *D. melanogaster* shows sexual size dimorphism, and that the sexes may vary in their plasticity (Hangartner et al., 2022; Pottier et al., 2021; Shingleton et al., 2017; Stillwell et al., 2010), we assessed size plasticity separately for males and females. By quantifying genetic variation in plasticity within a population in response to combinations of environmental stressors, our study sheds new insight into how animal populations may respond to ongoing climate change.

## Methods

### Fly stocks - isogenic lines

We used isogenic lines that were established by Chakraborty *et al*. (2023). Briefly, two hundred field-inseminated female *D. melanogaster* were collected in Duranbah, New South Wales, Australia (28.3° S, 153.5° E) in 2018. From each of these females, two independent iso-female lines were established and bred into four hundred lines. Full-sibling mating was performed in each of these four hundred lines for twenty generations following MacKay *et al*. (2012), resulting in 71 isogenic lines. Four copies of each isogenic line were reared at 25°C on standard diets (1:3 P:C ratio and 320 kcal/ L) under 12:12 light:dark cycle for twelve generations prior to the experiment, as described below. The standard diet was prepared by mixing water with potato flakes (18.18 g/ L), yeast (36.36 g/ L), agar (6.36 g/ L), and dextrose (27.27 g/ L), with addition of nipagin (6 mL/L) and propionic acid (2.5 mL/L) to prevent bacterial and fungal growth (Holleley et al., 2008). While we acknowledge the potential influence of gut microbiota on the life history traits examined, our study was designed to investigate the effects of combinations of nutrition and temperature within the constraints of our experimental design (described below); we therefore chose not to manipulate the gut microbiome so that the experimental diets (describe below) used in our study were treated the same. This meant that we could focus on the relative effects of nutrition without needing to also consider changes in the gut microbiome.

### Developmental treatment conditions and choice of traits

We used six thermal-nutritional conditions to simulate the possible shifts in temperature (Masson-Delmotte et al., 2018) and nutrition (DaMatta *et al*. 2010; Gowik and Westhoff 2011; IPCC 2018) under climate change. They included combinations of two temperatures: 25°C and 28°C, and three diets: standard diet (1:3 P:C ratio and 320 kcal/L), low caloric diet (1:3 P:C and 80kcal/L, a 25% dilution of the standard diet), and low P:C diet (1:12 P:C and 320kcal/L). 25°C represents the average summer temperature in south-east Australia, whilst 28°C represents the 3°C temperature increase projected in twenty years under climate change (Screen, 2014). The low caloric diet and the low P:C diets simulate a reduction of protein and variation of carbohydrates in plant composition projected under climate change (DaMatta *et al*. 2010). Importantly, they also reflect the nutritional range in the rotting fruit upon which *Drosophila* feed in nature (Matavelli et al., 2015; Silva-Soares et al., 2017). These conditions have been previously shown to successfully induce plastic responses in our traits of interest, egg-to-adult viability, and wing size (Chakraborty et al., 2020; Kutz et al., 2019).

Egg-to-adult viability and wing size were chosen because they are fitness-related developmental traits known to be sensitive to nutrition and diet individually and in combination (Chakraborty et al., 2020; Kutz et al., 2019). Egg-to-adult viability is a direct indication of an individual’s ability to survive to the reproductive adult stage (Kristensen et al., 2015). While viability data by its nature includes missing data as we are unable to measure the plasticity of traits such as body size in individuals that did not survive development (Hadfield, 2008), it nonetheless represents a trait linked to fitness that has previously been used in studies of plasticity (David et al., 2004; Kellermann & van Heerwaarden, 2019). Larger adult body size often correlates with greater fitness in ectotherms, including *Drosophila* (Kingsolver & Huey, 2008), and so plasticity in this trait will likely affect fitness.

### Experimental set up - Egg lays to obtain experimental flies

Adults from the same copy of the isogenic lines were mixed and transferred to laying plates containing apple juice, agar, nipagin, and a surface layer of autoclaved yeast to encourage oviposition. Laying plates were maintained at 25°C for 18 hours to allow egg laying.

### Egg-to-adult viability assay

Eggs were collected from laying plates and transferred to vials containing 7.2 ml of their respective diets. Three replicate vials were set up for each treatment and line combination, with a density of twenty eggs per vial (Agnew et al., 2002). Vials containing fresh eggs were kept at their respective treatment temperature at a 12:12 hour light: dark cycle until adult eclosion. Egg-to-adult viability was calculated as the number of eclosed adults relative to the initial number of eggs deposited into each vial (20 eggs). This resulted in a sample size of 71 isogenic lines x 2 temperatures x 3 diets x 3 replicate vials, which equals 1278 vials. Eggs were picked across 52 days, which are defined as block for the statistical analyses (described below).

### Adult body size measurement

Wing size was used as a proxy for body size (Partridge et al., 1994). After eclosing, adults were sexed and preserved in a 70% ethanol-30% glycerol solution for wing dissection. The left wings of each sex were removed with fine forceps, transferred to glass slides, and suspended in 70% ethanol-30% glycerol solution for imaging. To measure the wing centroid size, landmarks were positioned at eight vein intersections as indicated in Fig. 1, and the X-and Y-coordinates of landmarks were determined using the software package tpsDIG (Rohlf, 2015). Using the software CoorDGen8 (Sheets, 2014), wing size was estimated using centroid size, the square root of the sum of the squared distances between each landmark (Rohlf & Slice, 1990).

**Fig. 1.**
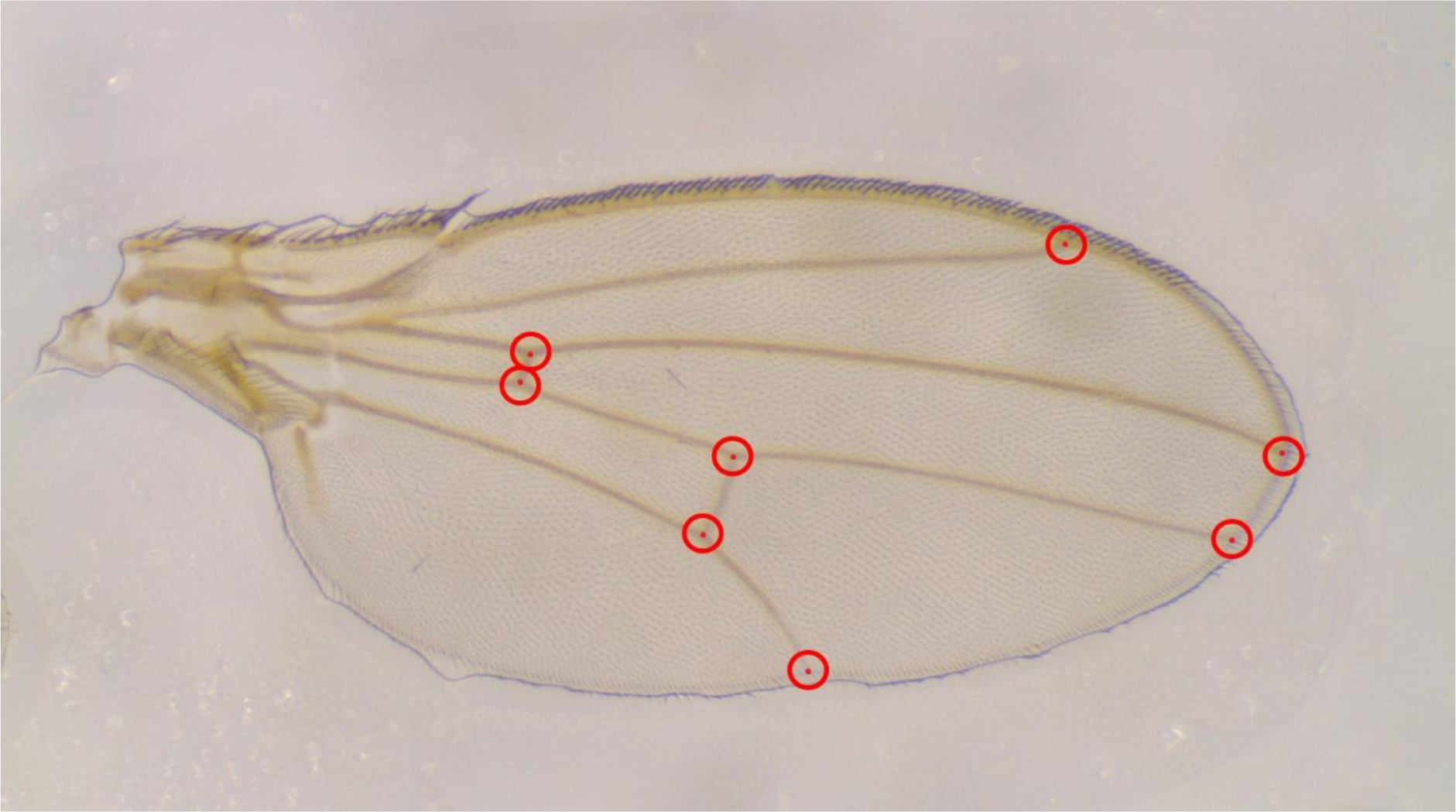
Landmarks used for calculating wing centroid size, indicted by red dots.

### Statistical analysis

#### Testing for overall genotype-by-environment (G × E) interactions

R was used to perform all data analysis and visualization (version 3.4.3) (R Core Team, 2017). To test for significant G × E, we used the *lme4* package (Bates et al., 2015) to fit linear mixed effects models for both traits. A binomial distribution was used to fit models for egg-to-adult viability, while wing size was log-transformed (Labarbera, 1989), fitted with a Gaussian error distribution, and analysed separately for each sex.

Models were fitted separately for each trait. Random effects of block (defined as the days of egg picking), line, temperature-by-line, diet-by-line, and temperature-by-diet-by-line were added sequentially (see equations 1-4) and the significance of each variable was tested with log-likelihood ratio tests using the *lrtest* package (Zeileis and Hothorn, 2002).

Specifically, the first model (equation 1) included temperature and diet as the fixed effects, and block as a random effect. This model estimates the overall population response to combinations of temperature and diet while accounting for the differences in traits among blocks. Vial is not included in any of the models as a random effect because initial analyses revealed it had no significant effect.

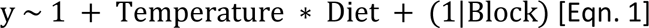

In the second model (equation 2), line was added as an additional random effect to test for differences in traits across the isogenic lines.

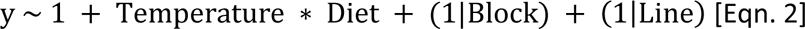

In the third model (equation 3), temperature and diet were added to the random line term to test for differences across lines in their plastic response in response to temperature and diet individually (i.e. G × E), allowing random intercepts and slopes.

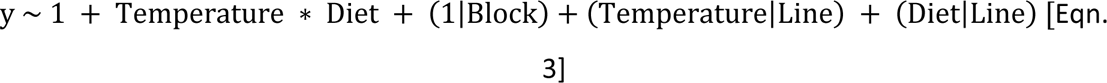

In the fourth model (equation 4), a temperature-by-diet-by-line interaction was further added as a random effect to test for differences across lines in their plastic responses to the combined effects of diet and temperature (i.e. overall G × E), allowing random intercepts. Comparing Eqn. 4 to Eqn. 3 tests whether plasticity in response to the combined stressors differed from the plastic response to individual stressors.

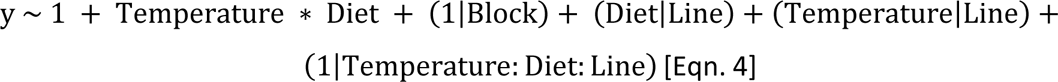

In the last model (equation 5), a temperature-by-diet term was added to the random line term. Comparing the fit of Eqn. 5 to Eqn. 3 tests for significant G × E as differences among lines in both intercepts and slopes (i.e. allowing changes in expression of genetic variance and correlations as the cause of G × E).

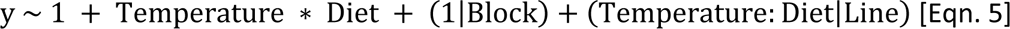

#### Quantifying genetic (co)variance within and across environments

Genetic (co)variances for egg-to-adult viability and wing size were estimated across treatments, and across sexes (the latter only for wing size) to illustrate the extent to which the observed G × E was due to changes in genetic variance across environments and/or genetic correlations that are less than one (Via & Lande, 1985). Because isogenic lines were used in this study, all estimates of genetic variance and genetic correlations are broad sense – they include additive and non-additive variance components.

Specifically, we used the *MCMCglmm* package (Hadfield, 2010) to fit models using Markov Chain Monte Carlo (MCMC), to estimate genetic (co)variances and genetic correlations. For each model, we used 1 million iterations with a burn-in of 100,000 iterations and a thinning interval of 500 iterations. We checked model convergence by ensuring that effective sample sizes were more than 85% of samples saved. We used weakly-informative parameter-expanded priors for the variance components, and checked their sensitivity by changing the scale of the parameters while ensuring there were no changes in the posterior distributions. A thousand iterations were kept from each model, providing the posterior distribution for all parameters estimated. In all estimations of genetic (co)variance, each combination of diet (standard, low calorie, and low P:C), temperature (25°C and 28°C), and sex (only for wing size) were treated as separate traits/treatments. Models were analysed using raw viability data and log-transformed wing size data.

#### Quantifying genetic (co)variance in egg-to-adult viability and wing size in each sex

To estimate genetic (co)variance in egg-to-adult viability within and across environments, we fitted viability data using a binomial distribution with the generalised linear mixed model.

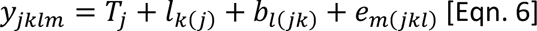

We included viability as the response variable *y_jklm_* and treatment *T_j_* as the only fixed effect. Random effects included isogenic line *l_k(j)_* and block *b_l(j)_* within the treatments, to remove any variance due to differences between experimental blocks. The term *e_m(jkl)_* represents the residuals. For the isogenic line component, we estimated an unstructured covariance matrix, which calculates the genetic variance of egg-to-adult viability within each treatment, and the genetic covariance between treatments, resulting in a 6 × 6 genetic (co)variance matrix. Genetic correlations between treatments were calculated from the (co)variance matrix.

Genetic (co)variances and genetic correlations for wing size within and across environments for each sex were estimated as described above using a Gaussian distribution.

#### Quantifying cross-sex genetic correlations in wing size

We analysed cross-sex correlations of wing size within- and between-treatments using this equation,

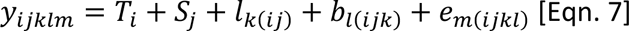

where sex *S_j_* was included as a fixed effect. For the isogenic line component, we estimated an unstructured covariance matrix that estimates the genetic variances of wing size among isogenic lines in each combination of treatment and sex, and genetic covariances between combinations respectively. We subset data by treatments and combinations of treatment and sex to reduce the number of (co)variance parameters estimated. Six 4 × 4 genetic (co)variance matrices were generated for calculating within-treatment cross-sex genetic correlations, and fifteen 2 × 2 genetic (co)variance matrices were generated for calculating between-treatment cross-sex genetic correlations.

#### Genetic correlations between egg-to-adult viability and wing size

To estimate genetic correlations between egg-to-adult viability and wing size across treatments, we applied equation 6 but included both egg-to-adult viability and wing size as a bivariate response variable (note that wing size was averaged over both sexes and estimated without sex as a fixed effect). As egg-to-adult viability and wing size were measured on different scales, we standardised each trait by dividing trait values by their grand mean (Houle, 1992). We applied equation 6 separately to the data from each treatment, and only included isogenic line and block as random effects. An unstructured covariance matrix was estimated, which calculates the genetic variance in viability and wing size, and the genetic covariance between viability and wing size within each treatment. This produced six 2 × 2 genetic (co)variance matrices. Genetic correlations between traits for each treatment were then calculated from each (co)variance matrix.

## Results

We characterised genetic variation in plasticity (G × E) for both egg-to-adult viability and sex-specific wing size in response to combined stressors within a population. We subjected *D. melanogaster* isogenic lines to six different thermal-nutritional conditions (25°C— standard diet, 25°C—low caloric diet, 25°C—low P:C diet, 28°C—standard diet, 28°C—low caloric diet, and 28°C—low P:C diet) and tested for significant genetic variation in plasticity in egg-to-adult viability and wing size. We then calculated genetic (co)variances and genetic correlations for each trait within and across environments and sexes (for wing size) to understand the causes of the significant G × E. Genetic correlations between egg-to-adult viability and wing size were also calculated to understand the independent evolvability between traits.

### Genotype-by-environment interactions in egg-to-adult viability

We found evidence of significant genetic variation in plasticity for viability (G × E) in response to temperature and diet individually. Importantly, the plastic response to combinations of temperature and nutrition varied from the plastic responses seen for either temperature or nutrition individually (Table 1). When we look at the raw data, such variation in plasticity is captured by non-parallel reaction norms among lines across diets and temperatures (Fig. 2A).

**Table 1.** Linear models testing for genotype (Line) by environment interactions in egg-to-adult viability in response to combined developmental temperature and diet conditions.

| Model number | Egg-to-adult viability (y) model | Likelihood ratio test | P-value |
| --- | --- | --- | --- |
| 1.1 | $y \sim 1 + \text{Temperature} * \text{Diet} + (1 \vert \text{Block})$ | - | - |
| 1.2 | $y \sim 1 + \text{Temperature} * \text{Diet} + (1 \vert \text{Block}) + (1 \vert \text{Line})$ | 1.2 to 1.1 | <0.001 |
| 1.3 | $y \sim 1 + \text{Temperature} * \text{Diet} + (1 \vert \text{Block}) + (\text{Temperature} \vert \text{Line}) + (\text{Diet} \vert \text{Line})$ | 1.3 to 1.2 | <0.001 |
| 1.4 | $y \sim 1 + \text{Temperature} * \text{Diet} + (1 \vert \text{Block}) + (\text{Temperature} \vert \text{Line}) + (\text{Diet} \vert \text{Line}) + (1 \vert \text{Temperature:Diet:Line})$ | 1.4 to 1.3 | <0.001 |
| 1.5 | $y \sim 1 + \text{Temperature} * \text{Diet} + (1 \vert \text{Block}) + (\text{Temperature:Diet} \vert \text{Line})$ | 1.5 to 1.3 | <0.001 |

**Fig. 2.**
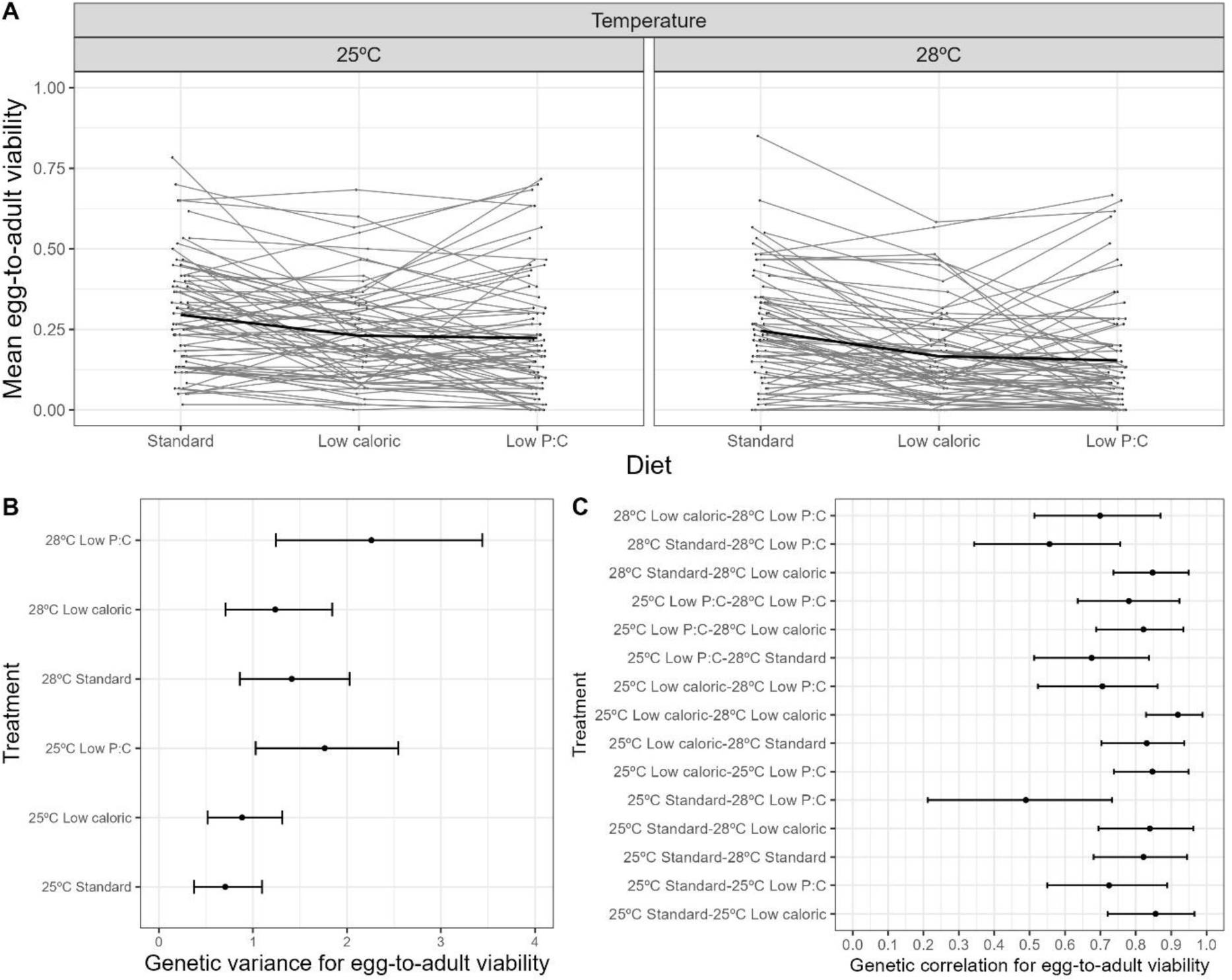
(**A**) Reaction norms of mean egg-to-adult viability of 71 isogenic lines across six thermal-nutritional treatments. Solid black line in both panels represents the overall population mean reaction norm, and the light grey lines the reaction norms for each isogenic line (genotype) across the three diets. For ease of visualisation we have separated the panels by temperature. (**B**) Estimates of genetic variance (which includes non-additive genetic variance) for egg-to-adult viability in response to six combined thermal-nutritional treatments, and (**C**) genetic correlation (broad sense) for egg-to-adult viability across treatments. Error bars show 95% upper and lower highest posterior density (HPD) intervals in panels B and C.

### Genetic (co)variances in egg-to-adult viability

We estimated genetic (co)variance matrices to determine whether significant G × E for viability was caused by differences in the amount of genetic variance across environments and/or cross-environment genetic correlations that are less than one (Via & Lande, 1985). We found significant genetic variance for viability in all treatments (Fig. 2B), and that genetic variance for viability at 28°C low P:C diet was significantly higher than the genetic variance at 25°C standard diet (Fig. 2B and Fig. S1), evidenced by the non-overlapping 95% highest posterior density (HPD) intervals (Fig. 2B).

Genetic correlations between treatment pairs were all positive and significantly greater than zero, but mostly less than one (Fig. 2C and Fig. S2). This suggests that significant G × E for viability results from both differences in the expression of genetic variance across environments, and differences in how isogenic lines respond to environments (Falconer, 1952; Via & Lande, 1985).

### Genotype-by-environment interactions in wing size

We found evidence for genetic variation in plasticity for wing size in males and females in response to temperature and nutrition individually, and that the plastic response of size to combinations of temperature and nutrition varied from the plastic responses seen for temperature and nutrition individually (Table 2). Such significant within-population G × E in male and female wing size is visualised as non-parallel reaction norms in Fig. 3A-B.

**Table 2.** Linear models testing for genotype (Line) by environment interactions in wing size of both sexes in response to combined developmental temperature and diet conditions.

| Model number | Male wing size (y) model | Likelihood ratio test | P-value |
| --- | --- | --- | --- |
| 2.1 | $\log(y) \sim 1 + \text{Temperature} * \text{Diet} + (1 \vert \text{Block})$ | - | - |
| 2.2 | $\log(y) \sim 1 + \text{Temperature} * \text{Diet} + (1 \vert \text{Block}) + (1 \vert \text{Line})$ | 2.2 to 2.1 | <0.001 |
| 2.3 | $\log(y) \sim 1 + \text{Temperature} * \text{Diet} + (1 \vert \text{Block}) + (\text{Temperature} \vert \text{Line}) + (\text{Diet} \vert \text{Line})$ | 2.3 to 2.2 | <0.001 |
| 2.4 | $\log(y) \sim 1 + \text{Temperature} * \text{Diet} + (1 \vert \text{Block}) + (\text{Temperature} \vert \text{Line}) + (\text{Diet} \vert \text{Line}) + (1 \vert \text{Temperature:Diet:Line})$ | 2.4 to 2.3 | <0.001 |
| 2.5 | $\log(y) \sim 1 + \text{Temperature} * \text{Diet} + (1 \vert \text{Block}) + (\text{Temperature:Diet} \vert \text{Line})$ | 2.5 to 2.3 | 0.007 |
| **Female wing size (y) model** |  |  |  |
| 3.1 | $\log(y) \sim 1 + \text{Temperature} * \text{Diet} + (1 \vert \text{Block})$ | - | - |
| 3.2 | $\log(y) \sim 1 + \text{Temperature} * \text{Diet} + (1 \vert \text{Block}) + (1 \vert \text{Line})$ | 3.2 to 3.1 | <0.001 |
| 3.3 | $\log(y) \sim 1 + \text{Temperature} * \text{Diet} + (1 \vert \text{Block}) + (\text{Temperature} \vert \text{Line}) + (\text{Diet} \vert \text{Line})$ | 3.3 to 3.2 | <0.001 |
| 3.4 | $\log(y) \sim 1 + \text{Temperature} * \text{Diet} + (1 \vert \text{Block}) + (\text{Temperature} \vert \text{Line}) + (\text{Diet} \vert \text{Line}) + (1 \vert \text{Temperature:Diet:Line})$ | 3.4 to 3.3 | <0.001 |
| 3.5 | $\log(y) \sim 1 + \text{Temperature} * \text{Diet} + (1 \vert \text{Block}) + (\text{Temperature:Diet} \vert \text{Line})$ | 3.5 to 3.3 | <0.001 |

**Fig. 3.**
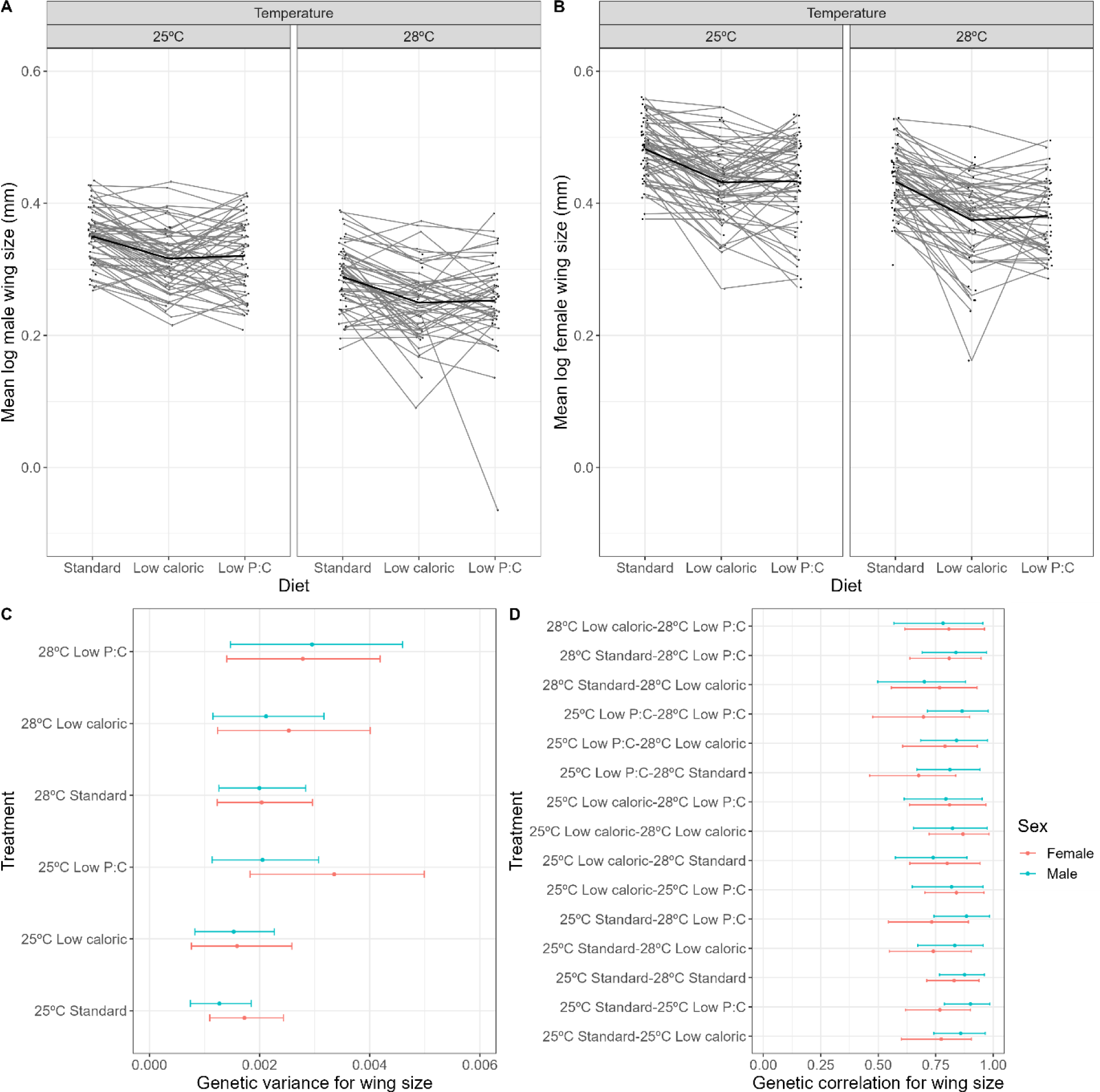
Reaction norms of mean log wing size of (**A**) males and (**B**) females of 71 isogenic lines across six combined thermal-nutritional treatments. Solid black line in each panel represents the overall population mean reaction norm, and the light grey lines the reaction norms for each isogenic line (genotype) across the three diets. For ease of visualisation we have separated the panels by temperature. (**C**) Estimates of genetic variance (which includes non-additive genetic variance) for males and female wing size in response to six combine thermal-nutritional treatments, and (**D**) Genetic correlation (broad sense) for wing size across treatments for males and females. Error bars show 95% upper and lower HPD intervals in panels C and D.

### Genetic (co)variances in wing size

We found significant genetic variance in wing size for all treatments within each sex; however, the distribution of genetic variance did not differ across treatments (Fig. 3C and Fig. S3). Thus, significant G × E in wing size in both males and females was not caused by the differences in the amount of genetic variance across the environments.

Genetic correlations in wing size across the six thermal-nutritional treatments within each sex were all positive, significantly greater than zero, and mostly less than one (Fig. 3D, Fig. S4 and Fig. S5). This suggests that significant G × E in wing size of both males and females was caused by differences in the alleles contributing to the response across environments (Falconer, 1952; Via & Lande, 1985).

### Cross-sex genetic correlations for wing size

To determine whether wing size has a similar genetic basis in both sexes, we estimated the cross-sex genetic correlations for wing size within and between all thermal-nutritional treatments. This helps us to understand the potential for size to evolve independently in each sex and in each environment (Falconer, 1952; Via & Lande, 1985).

Cross-sex correlations within treatments (indicated in red in Fig. 4, Fig. S6) were significantly greater than zero, and large, but less than one, indicating that even though sexes share some genetic basis for these traits, there is also some degree of genetic independence. Some cross-sex genetic correlations were significantly smaller when males and females developed in different treatments (i.e. male: 25°C low P:C – female: 28°C Standard; male: 28°C standard – female: 25°C low caloric, 25°C low P:C, and 28°C low caloric; male: 28°C low caloric – female: 25°C standard and 28°C standard; indicated in blue in Fig. 4). This suggests that the genetic basis for wing size in males and females diverges when the sexes are reared in different environments, which could allow sexes to evolve independently as conditions become more variable.

**Fig. 4.**
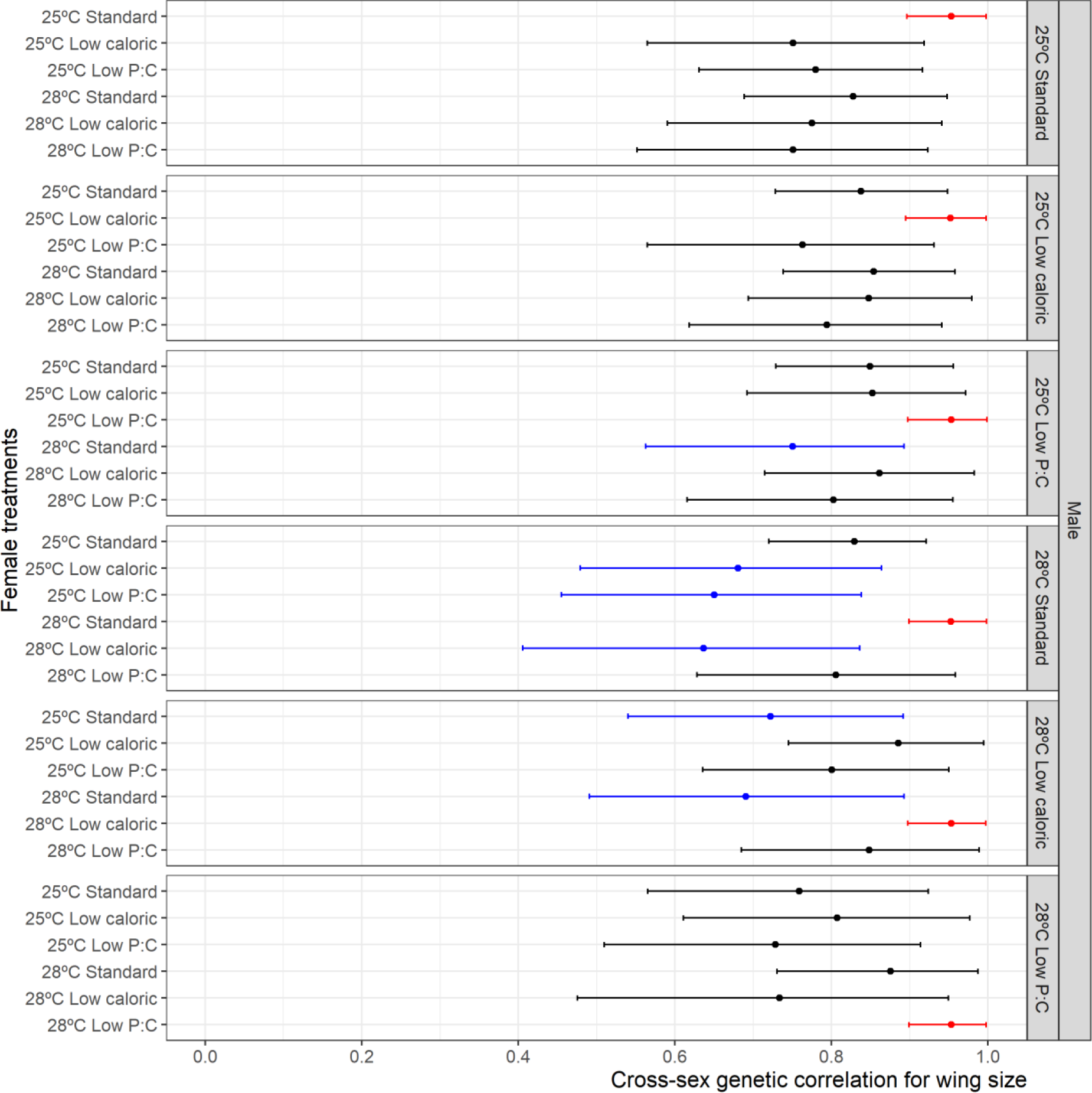
Cross-sex genetic correlation (broad sense) for wing size across six combined thermal-nutritional treatments. Error bars show 95% upper and lower HPD intervals. Red error bars indicate the cross-sex genetic correlation within treatments. The blue error bars show significantly smaller cross-sex genetic correlations across certain treatment combinations compared to the cross-sex genetic correlations within treatments (red error bars).

### Genetic correlations between egg-to-adult viability and wing size

Finally, we estimated genetic correlations between egg-to-adult viability and wing size (sexes combined) across all thermal-nutritional treatments to determine whether these traits have a similar genetic basis, and whether they can evolve independently in different environments (Falconer, 1952; Via & Lande, 1985). Correlations between viability and wing size were not significantly different from zero in all treatments (Fig. 5 and Fig. S7), indicating that these traits are genetically independent.

**Fig. 5.**
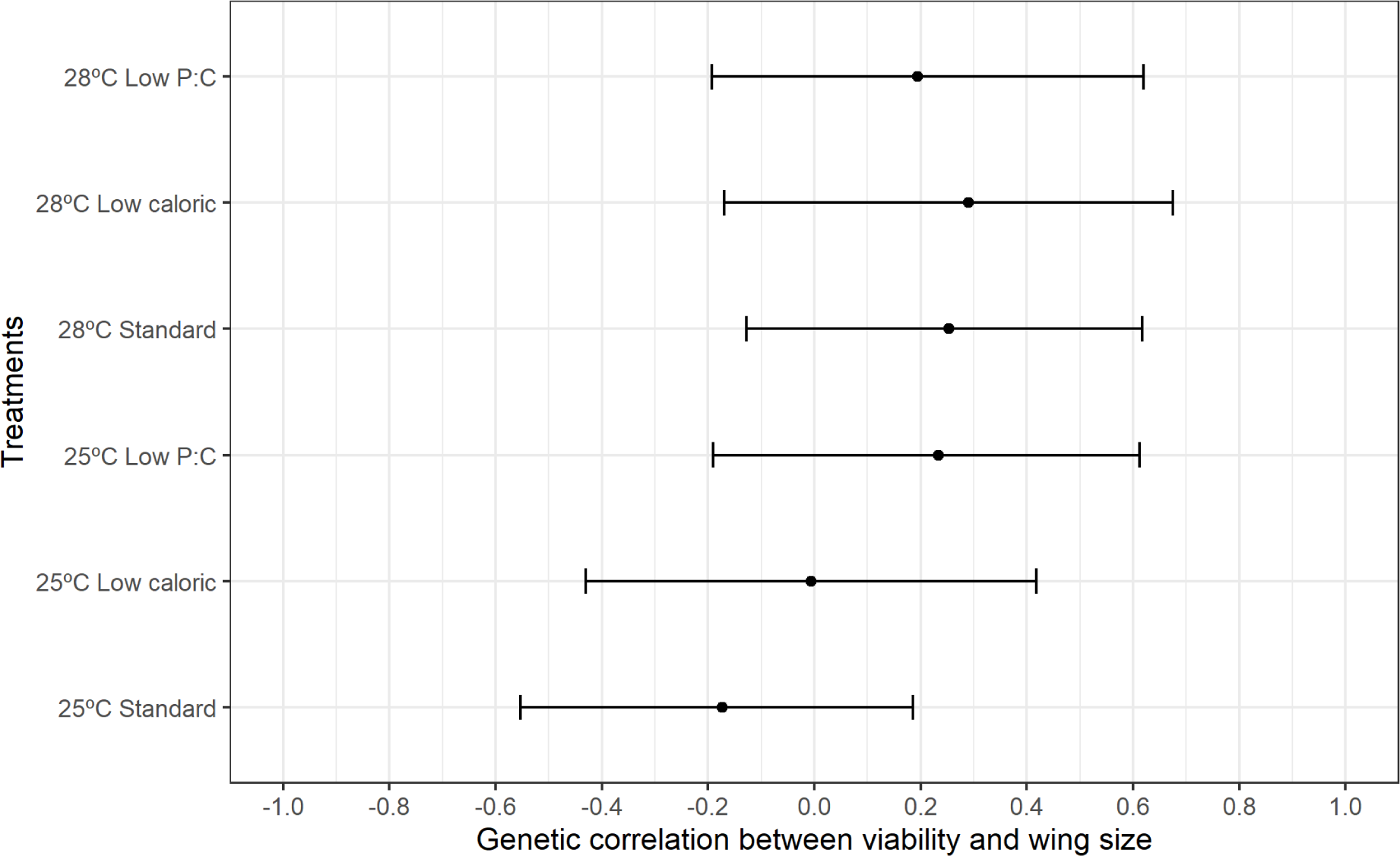
Genetic correlation (broad sense) between egg-to-adult viability and wing size within each of six combined thermal-nutritional treatments. Wing size of both sexes was combined for estimating the genetic correlation between egg-to-adult viability and wing size. Error bars shows 95% upper and lower HPD intervals.

## Discussion

Climate change is simultaneously imposing thermal and nutritional stress on animals (Pecl et al., 2017; Van Der Putten et al., 2010), and is expected to negatively affect biodiversity in the long term (Masson-Delmotte et al., 2018; Rosenblatt & Schmitz, 2016; Walters et al., 2012). Phenotypic plasticity is an important means by which animals will mitigate the increasing stress under climate change (Seebacher et al., 2015; Sgrò et al., 2016; Urban et al., 2014), although it may not always be adaptive (Ghalambor et al., 2007; Sgrò et al., 2016). Plastic responses of life-history traits can shift in response to changes in temperature (French et al., 1998; Hoffmann et al., 2003; Schou et al., 2017) and nutrition individually (Bader & Williams, 2012; Jang & Lee, 2018; Min et al., 2007). Temperature can also modify plastic responses of life history traits to nutritional stress, causing an interactive effect (Chakraborty et al., 2020; Kutz et al., 2019; Lee & Roh, 2010). While these studies improve our understanding of how animals may respond to combined stress resulting from climate change, they have not accounted for genetic variation in plasticity within a population.

Individuals within a population can show substantial variation in plasticity (Chakraborty et al., 2023; Frankino et al., 2019; Lafuente et al., 2018; Mossman et al., 2016; J. B. Wang et al., 2017). Understanding within-population variation matters because it may shape the capacity for populations to adapt and persist in the face of ongoing environmental change (Chevin et al., 2010; Dreyer et al., 2016; Frankino et al., 2019; Lande, 2009). Extensive within-population variation in plasticity has been reported in a range of life-history traits such as viability (Duun Rohde et al., 2016) and body size (Barker & Krebs, 1995; Imasheva et al., 1999; Lafuente et al., 2018) in response to thermal and nutritional stress individually (Frankino et al., 2019; Lafuente et al., 2018; Mossman et al., 2016; J. B. Wang et al., 2017). The results of the current study are consistent with these findings. Importantly, we also show that within-population variation in viability and body size plasticity in response to combinations of temperature and diet differed from the plastic responses to temperature and nutrition individually.

The extent to which genotypes within a population vary in their plasticity will determine how the population responds to selection imposed by environmental change (Chevin et al., 2010; Dreyer et al., 2016; Frankino et al., 2019). Populations with similar mean plasticity (reaction norms) may vary markedly in their genotype-specific reaction norms, causing them to vary in their response to selection (Dreyer et al., 2016; Frankino et al., 2019; Lande, 2009). Predictions of long-term evolutionary responses to environmental changes using population-level assessments of plasticity, could therefore be inaccurate unless within-population variation in plasticity is also considered (Dreyer et al., 2016; Frankino et al., 2019; Reed et al., 2010).

We found that the significant G × E for viability was the result of both significant differences in the expression of genetic variance across environments, and cross-environment genetic correlations that were greater than zero but less than one. Specifically, we found that genetic variance for viability was significantly higher at 28°C on a low P:C diet compared to 25°C on a standard diet. Bubliy *et al*. (2001) also observed that significant G × E in viability was due to an increase in genetic variance for viability at elevated (32°C) compared to control (25°C) temperatures. In contrast, Bubliy and Loeschcke (2002) found no differences in the expression of genetic variance for viability between 13°C and 25°C. The increase in genetic variance at elevated temperatures individually (Bubliy et al., 2001) or when combined with nutritional stress (current study) are consistent with the idea that higher levels of environmental stress can result in increases in the expression of genetic variance (Hoffmann & Merilä, 1999; Imasheva et al., 1998), although the mechanisms of such changes remains unknown. Our results suggest that the adaptive capacity of populations, at least with respect to viability, may be enhanced by some of the changes expected to occur with climate change, namely elevated temperature combined with diets low in protein but high in carbohydrates (Burla & Taylor, 1982; Fowler & Whitlock, 2002).

We found that the significant G × E interactions for viability were also the result of genetic correlations across environments that were above zero and often less than one. This indicates that viability across some thermal-nutritional conditions is controlled by different alleles or differently by the same alleles (Via & Lande, 1985), meaning that viability has some capacity to evolve independently across different thermal-nutritional environments (Via & Lande, 1987). While some studies have found that genetic correlations can change in sign and magnitude across environments (Sgrò & Hoffmann, 2004; Wood & Brodie, 2015), we did not find this to be the case for viability in the present study.

Although body size plasticity has been shown to vary within a population in response to temperature (Barker & Krebs, 1995; Cavicchi et al., 1995; Lafuente et al., 2018) and nutrition individually (Imasheva et al., 1999) and in combination (De Moed et al., 1997), the source of this G × E has differed across studies. De Moed *et al*. (1997) found that the significant G × E for body size was due to differences in the expression of genetic variance across environments; namely, the expression of genetic variance for size increased only under combinations of low temperature (15°C) and poor food (low protein). We found that the expression of genetic variance for size did not differ across environments in either sex; rather the significant G × E for body size for both sexes was due to genetic correlations across treatments that were significantly more than zero but often less than one. Similarly, Barker and Krebs (1995) and Imasheva *et al*. (1999) also found that genetic correlations less than unity, rather than changes in the expression of genetic variance across environments, were responsible for G × E for body size in response to temperature and dietary yeast concentration respectively. Interestingly, De Moed *et al*. (1997) did not find an increase in genetic variance for size when larvae developed at higher temperature (27.7 °C) and poor food, which is consistent with the present study.

Differences across studies in the extent to which genetic variance and/or genetic correlations change with environment may reflect the use of different source/experimental populations, environmental conditions, and experimental approaches across studies. Yet, studies to date (reviewed in Hoffmann and Merilä (1999); Sgrò and Hoffmann (2004); Wood and Brodie (2015)) indicate that although the environmental dependency of genetic variance and correlations may be common, is not easily predicted by trait or environment.

As expected from previous studies (Cowley et al., 1986; Cowley & Atchley, 1990; Lasne et al., 2018; Reeve & Fairbairn, 1996), we found strong and positive cross-sex genetic correlations for wing size within treatments. This indicates that the genetic basis of plasticity in wing size is largely shared between males and females, but because these correlations are less than one, there is some potential for size to evolve independently between sexes (Lande, 1980; Via & Lande, 1987). More importantly, we found that some cross-sex genetic correlations were reduced when males and females developed in different environments. This suggests that there is a greater possibility for the independent evolution of wing size when the sexes experience markedly different thermal-nutritional environments.

Most studies find that cross-sex genetic correlations are large and positive (Poissant et al., 2010). However, there is some evidence that cross-sex genetic correlations can change in magnitude (Delcourt et al., 2009) and sign (Berger et al., 2014). Specifically, in the seed beetle, the cross-sex genetic correlation for fitness changed from negative under benign temperatures to positive under thermal stress (Berger et al., 2014), while in *D. serrata*, Delcourt *et al*. (2009) showed that the negative cross-sex genetic correlation for fitness became stronger on a novel diet. In the wild, individual *Drosophila* are often found in different microhabitats – different temperatures due to daily thermal fluctuations (Noer et al., 2020) and different nutritional contexts due to different niches, resources (Diepenbrock & Burrack, 2017), and nutritional requirements (Camus et al., 2018; Duxbury & Chapman, 2021; Lee et al., 2013). Thus, weaker cross-sex genetic correlations under varying environmental conditions may facilitate adaptation if changing environments favour sex-specific optima, which is often the case (Berger et al., 2014; Delcourt et al., 2009; Poissant et al., 2010).

Finally, genetic correlations between viability and wing size across all six thermal-nutritional treatments were not different from zero. This means that these two traits do not share a similar genetic basis, and are able to evolve independently (Via & Lande, 1985). Our findings differ from earlier studies. For instance, Teuschl *et al*. (2007) artificially selected dung flies with different body sizes − small, control, and large – and assessed their pre-adult viability in combinations of high temperature and limited food. Interestingly, larger males had higher pre-adult mortality, suggesting body size and viability might be negatively genetically correlated. Dingle (1992) also observed a similar phenomenon in milkweed bugs under food shortage, such that higher pre-adult survival was traded off with smaller body size, which again indicates a negative genetic correlation between these two traits (Via & Lande, 1985). The fact that we found no correlation between viability and wing size suggests that the genetic correlation between viability and body size may be species- and condition-specific.

While this study shows that plastic responses to temperature and diet individually vary markedly from plastic shifts in response to temperature and diet in combination, we acknowledge that our experimental conditions are not a direct reflection of conditions experienced in nature. In particular, we used constant temperatures, when in nature temperatures fluctuate. Future studies should incorporate thermal variation and fluctuations to better reflect daily temperature fluctuations, since plastic responses of traits such as body size traits (Czarnoleski et al., 2013), fertility and productivity traits (Rodrigues et al., 2022) to varying temperature regimes can differ from those in response to constant temperatures. That said, our study nonetheless illustrates for the first time how plasticity in response to single stressors does not predict plastic shifts in response to the same stressors individually.

Finally, our estimates of genetic (co)variance include non-additive components of variation. In particular, epistatic variance can be significantly increased in highly inbred isogenic lines (Huang et al., 2020; Mackay, 2010). While positive and non-directional epistasis generally align with predictions from additive genetic variance, negative epistasis may lead to canalisation (Carter et al., 2005). Due to the potential contribution of epistatic variance to our estimates of genetic (co)variance, direct predictions of evolutionary responses to combinations of stressors from our data are not possible. Nonetheless, our data do allow us to quantify genetic variation for plasticity within a population across traits and sexes, and explore how the expression of genetic variation and covariation changes with the environment and sex.

In summary, we explicitly assessed the genetic basis of within-population variation in plasticity of egg-to-adult viability and wing size in *D. melanogaster* in response to combined thermal-nutritional conditions. We demonstrated that individual genotypes varied in their plastic responses, and that the plastic response to temperature and diet individually differed from the plastic response to both temperature and nutrition in combination. We also found that the source of G × E resulted from changes in the expression of genetic variance (viability only), and genetic correlations less than unity across environments (both traits). Additionally, our results suggest that the independent evolution of wing size between the sexes may be possible when the sexes experience different environmental conditions, and that viability and wing size are genetically independent, at least under the conditions assessed here. Our research sheds light on the complex genetic basis of life-history traits in response to combined stressors, and the potential for the genetic basis of traits to change across environments. Accurate predictions of how species may respond to climate change will require studies of plasticity that examine the longer-term consequences of G × E and sex-specific adaptation on evolutionary trajectories (Wood & Brodie, 2015).

## Bibliography

Abram, P. K., Boivin, G., Moiroux, J., & Brodeur, J. (2017). Behavioural effects of temperature on ectothermic animals: unifying thermal physiology and behavioural plasticity. Biological Reviews, 92(4), 1859–1876. 10.1111/brv.12312

Agnew, P., Hide, M., Sidobre, C., & Michalakis, Y. (2002). A minimalist approach to the effects of density-dependent competition on insect life-history traits. Ecological Entomology, 27(4), 396–402. 10.1046/j.1365-2311.2002.00430.x

Bader, C. A., & Williams, C. R. (2012). Mating, ovariole number and sperm production of the dengue vector mosquito Aedes aegypti (L.) in Australia: Broad thermal optima provide the capacity for survival in a changing climate. Physiological Entomology, 37(2), 136–144. 10.1111/j.1365-3032.2011.00818.x

Barker, J. S. F., & Krebs, R. A. (1995). Genetic variation and plasticity of thorax length and wing length in Drosophila aldrichi and D. buzzatii. Journal of Evolutionary Biology, 8(6), 689–709. 10.1046/j.1420-9101.1995.8060689.x

Bates, D., Mächler, M., Bolker, B. M., & Walker, S. C. (2015). Fitting linear mixed-effects models using lme4. Journal of Statistical Software, 67(1). 10.18637/jss.v067.i01

Berger, D., Grieshop, K., Lind, M. I., Goenaga, J., Maklakov, A. A., & Arnqvist, G. (2014). Intralocus sexual conflict and environmental stress. Evolution, 68(8), 2184–2196. 10.1111/evo.12439

Bochdanovits, Z., & De Jong, G. (2003). Experimental evolution in Drosophila melanogaster: Interaction of temperature and food quality selection regimes. Evolution, 57(8), 1829– 1836. 10.1111/j.0014-3820.2003.tb00590.x

Bowes, G. (1993). Facing the inevitable: Plants and increasing atmospheric CO2. Annual Review of Plant Physiology and Plant Molecular Biology, 44(1), 309–332. 10.1146/annurev.pp.44.060193.001521

Bubliy, O. A., Kristensen, T. N., Kellermann, V., & Loeschcke, V. (2012). Plastic responses to four environmental stresses and cross-resistance in a laboratory population of Drosophila melanogaster. Functional Ecology, 26(1), 245–253. 10.1111/j.1365-2435.2011.01928.x

Bubliy, O. A., & Loeschcke, V. (2002). Effect of low stressful temperature on genetic variation of five quantitative traits in Drosophila melanogaster. Heredity, 89(1), 70–75. 10.1038/sj.hdy.6800104

Bubliy, O. A., Loeschcke, V., & Imasheva, A. G. (2001). Genetic variation of morphological traits in Drosophila melanogaster under poor nutrition: isofemale lines and offspring-parent regression. Heredity, 86(3), 363–369. 10.1046/j.1365-2540.2001.00837.x

Burla, H., & Taylor, C. E. (1982). Increase of phenotypic variance in stressful environments. Journal of Heredity, 73(2), 142–142. 10.1093/oxfordjournals.jhered.a109599

Camus, M. F., Huang, C. C., Reuter, M., & Fowler, K. (2018). Dietary choices are influenced by genotype, mating status, and sex in Drosophila melanogaster. Ecology and Evolution, 8(11), 5385–5393. 10.1002/ece3.4055

Carter, A. J. R., Hermisson, J., & Hansen, T. F. (2005). The role of epistatic gene interactions in the response to selection and the evolution of evolvability. Theoretical Population Biology, 68(3), 179–196. 10.1016/j.tpb.2005.05.002

Cavicchi, S., Guerra, D., Torre, V. La, & Huey, R. B. (1995). Chromosomal Analysis of Heat-Shock Tolerance in Drosophila melanogaster Evolving at Different Temperatures in the Laboratory. Evolution, 49(4), 676. 10.2307/2410321

Chakraborty, A., Sgrò, C. M., & Mirth, C. K. (2020). Does local adaptation along a latitudinal cline shape plastic responses to combined thermal and nutritional stress? Evolution, 74(9), 2073–2087. 10.1111/evo.14065

Chakraborty, A., Walter, G. M., Monro, K., Alves, A. N., Mirth, C. K., & Sgrò, C. M. (2023). Within-population variation in body size plasticity in response to combined nutritional and thermal stress is partially independent from variation in development time. Journal of Evolutionary Biology, 36(1), 264–279. 10.1111/jeb.14099

Chevin, L. M., & Hoffmann, A. A. (2017). Evolution of phenotypic plasticity in extreme environments. Philosophical Transactions of the Royal Society B: Biological Sciences, 372(1723). 10.1098/rstb.2016.0138

Chevin, L. M., Lande, R., & Mace, G. M. (2010). Adaptation, plasticity, and extinction in a changing environment: Towards a predictive theory. PLoS Biology, 8(4). 10.1371/journal.pbio.1000357

Clissold, F. J., & Simpson, S. J. (2015). Temperature, food quality and life history traits of herbivorous insects. Current Opinion in Insect Science, 11, 63–70. 10.1016/j.cois.2015.10.011

Cockerell, F. E., Sgrò, C. M., & McKechnie, S. W. (2014). Latitudinal clines in heat tolerance, protein synthesis rate and transcript level of a candidate gene in Drosophila melanogaster. Journal of Insect Physiology, 60(1), 136–144. 10.1016/j.jinsphys.2013.12.003

Couret, J., Dotson, E., & Benedict, M. Q. (2014). Temperature, larval diet, and density effects on development rate and survival of Aedes aegypti (Diptera: Culicidae). PLoS ONE, 9(2). 10.1371/journal.pone.0087468

Cowley, D. E., & Atchley, W. R. (1990). Development and quantitative genetics of correlation structure among body parts of Drosophila melanogaster. The American Naturalist, 135(2), 242–268. 10.1086/285041

Cowley, D. E., Atchley, W. R., & Rutledge, J. J. (1986). Quantitative Genetics of Drosophila Melanogaster. I. Sexual Dimorphism in Genetic Parameters for Wing Traits. Genetics, 114(2), 549–566. 10.1093/genetics/114.2.549

Czarnoleski, M., Cooper, B. S., Kierat, J., & Angilletta, M. J. (2013). Flies developed small bodies and small cells in warm and in thermally fluctuating environments. Journal of Experimental Biology, 216(15), 2896–2901. 10.1242/jeb.083535

DaMatta, F. M., Grandis, A., Arenque, B. C., & Buckeridge, M. S. (2010). Impacts of climate changes on crop physiology and food quality. Food Research International, 43(7), 1814– 1823. 10.1016/j.foodres.2009.11.001

David, J. R., Allemand, R., Capy, P., Chakir, M., Gibert, P., Pétavy, G., & Moreteau, B. (2004). Comparative life histories and ecophysiology of Drosophila melanogaster and D. simulans. Genetica, 120(1–3), 151–163. 10.1023/B:GENE.0000017638.02813.5a

De Moed, G. H., De Jong, G., & Scharloo, W. (1997). Environmental effects on body size variation in Drosophila melanogaster and its cellular basis. Genetical Research, 70(1), 35–43. 10.1017/S0016672397002930

Delcourt, M., Blows, M. W., & Rundle, H. D. (2009). Sexually antagonistic genetic variance for fitness in an ancestral and a novel environment. Proceedings of the Royal Society B: Biological Sciences, 276(1664), 2009–2014. 10.1098/rspb.2008.1459

Diepenbrock, L. M., & Burrack, H. J. (2017). Variation of within-crop microhabitat use by Drosophila suzukii (Diptera: Drosophilidae) in blackberry. Journal of Applied Entomology, 141(1–2), 1–7. 10.1111/jen.12335

Diffenbaugh, N. S., & Field, C. B. (2013). Changes in ecologically critical terrestrial climate conditions. Science, 341(6145), 486–492. 10.1126/science.1237123

Dingle, H. (1992). Food level reaction norms in size-selected milkweed bugs Oncopeltus fasciatus. Ecological Entomology, 17(2), 121–126. 10.1111/j.1365-2311.1992.tb01168.x

Dreyer, A. P., Saleh Ziabari, O., Swanson, E. M., Chawla, A., Frankino, W. A., & Shingleton, A. W. (2016). Cryptic individual scaling relationships and the evolution of morphological scaling. Evolution, 70(8), 1703–1716. 10.1111/evo.12984

Duun Rohde, P., Krag, K., Loeschcke, V., Overgaard, J., Sørensen, P., & Nygaard Kristensen, T. (2016). A quantitative genomic approach for analysis of fitness and stress related traits in a Drosophila melanogaster model population. International Journal of Genomics, 2016(Ld). 10.1155/2016/2157494

Duxbury, E. M. L., & Chapman, T. (2021). Sex-specific responses of life span and fitness to variation in developmental versus adult diets in drosophila melanogaster. Journals of Gerontology - Series A Biological Sciences and Medical Sciences, 75(8), 1431–1438. 10.1093/GERONA/GLZ175

Dwyer, S. A., Ghannoum, O., Nicotra, A., & Von Caemmerer, S. (2007). High temperature acclimation of C4 photosynthesis is linked to changes in photosynthetic biochemistry. *Plant*, Cell and Environment, 30(1), 53–66. 10.1111/j.1365-3040.2006.01605.x

Economos, A. C., & Lints, F. A. (1984). Growth rate and life span in Drosophila. I. Methods and mechanisms of variation of growth rate. Mechanisms of Ageing and Development, 27(1), 1–13. 10.1016/0047-6374(84)90078-2

Falconer, D. S. (1952). The problem of environment and selection. The American Naturalist, 86(830), 293–298. 10.1086/281736

Fowler, K., & Whitlock, M. C. (2002). Environmental stress, inbreeding, and the nature of phenotypic and genetic variance in Drosophila melanogaster. Proceedings of the Royal Society B: Biological Sciences, 269(1492), 677–683. 10.1098/rspb.2001.1931

Franěk, R., Baloch, A. R., Kašpar, V., Saito, T., Fujimoto, T., Arai, K., & Pšenička, M. (2020). Isogenic lines in fish – a critical review. Reviews in Aquaculture, 12(3), 1412–1434. 10.1111/raq.12389

Frankino, W. A., Bakota, E., Dworkin, I., Wilkinson, G. S., Wolf, J. B., & Shingleton, A. W. (2019). Individual cryptic scaling relationships and the evolution of animal form. Integrative and Comparative Biology, 59(5), 1411–1428. 10.1093/icb/icz135

French, V., Feast, M., & Partridge, L. (1998). Body size and cell size in Drosophila: The developmental response to temperature. Journal of Insect Physiology, 44(11), 1081– 1089. 10.1016/S0022-1910(98)00061-4

Ghalambor, C. K., McKay, J. K., Carroll, S. P., & Reznick, D. N. (2007). Adaptive versus non-adaptive phenotypic plasticity and the potential for contemporary adaptation in new environments. Functional Ecology, 21(3), 394–407. 10.1111/j.1365-2435.2007.01283.x

Gowik, U., & Westhoff, P. (2011). The Path from C3 to C4 photosynthesis. Plant Physiology, 155(1), 56–63. 10.1104/pp.110.165308

Hadfield, J. D. (2008). Estimating evolutionary parameters when viability selection is operating. Proceedings of the Royal Society B: Biological Sciences, 275(1635), 723–734. 10.1098/rspb.2007.1013

Hangartner, S., Sgrò, C. M., Connallon, T., & Booksmythe, I. (2022). Sexual dimorphism in phenotypic plasticity and persistence under environmental change: An extension of theory and meta-analysis of current data. *Ecology Letters*, December 2021, 1–16. 10.1111/ele.14005

Hawlena, D., & Schmitz, O. J. (2010). Herbivore physiological response to predation risk and implications for ecosystem nutrient dynamics. Proceedings of the National Academy of Sciences of the United States of America, 107(35), 15503–15507. 10.1073/pnas.1009300107

Hoffmann, A. A., & Merilä, J. (1999). Heritable variation and evolution under favourable and unfavourable conditions. Trends in Ecology and Evolution, 14(3), 96–101. 10.1016/S0169-5347(99)01595-5

Hoffmann, A. A., Sørensen, J. G., & Loeschcke, V. (2003). Adaptation of Drosophila to temperature extremes: Bringing together quantitative and molecular approaches. Journal of Thermal Biology, 28(3), 175–216. 10.1016/S0306-4565(02)00057-8

Holleley, C. E., Hocking, A. D., Schubert, T. L., & Whitehead, M. R. (2008). Control of Penicillium roqueforti (Thom) infection in cultures of Drosophila melanogaster (Meigen) (Diptera: Drosophilidae). Australian Journal of Entomology, 47(2), 149–152. 10.1111/j.1440-6055.2007.00630.x

Houle, D. (1992). Comparing evolvability and variability of quantitative traits. Genetics, 130(1), 195–204. 10.1093/genetics/130.1.195

Huang, W., Carbone, M. A., Lyman, R. F., Anholt, R. R. H., & Mackay, T. F. C. (2020). Genotype by environment interaction for gene expression in Drosophila melanogaster. Nature Communications, 11(1), 1–10. 10.1038/s41467-020-19131-y

Huey, R. B., Carlson, M., Crozier, L., Frazier, M., Hamilton, H., Harley, C., Hoang, A., & Kingsolver, J. G. (2002). Plants versus animals: Do they deal with stress in different ways? Integrative and Comparative Biology, 42(3), 415–423. 10.1093/icb/42.3.415

Imasheva, A. G., Bosenko, D. V., & Bubli, O. A. (1999). Variation in morphological traits of Drosophila melanogaster (fruit fly) under nutritional stress. Heredity, 82(2), 187–192. 10.1038/sj.hdy.6884660

Imasheva, A. G., Loeschcke, V., Zhivotovsky, L. A., & Lazebny, O. E. (1998). Stress temperatures and quantitative variation in Drosophila melanogaster. Heredity, 81(3), 246–253. 10.1038/sj.hdy.6883840

Jang, T., & Lee, K. P. (2018). Comparing the impacts of macronutrients on life-history traits in larval and adult Drosophila melanogaster: The use of nutritional geometry and chemically defined diets. Journal of Experimental Biology, 221(21). 10.1242/jeb.181115

Janowitz, S. A., & Fischer, K. (2011). Opposing effects of heat stress on male versus female reproductive success in Bicyclus anynana butterflies. Journal of Thermal Biology, 36(5), 283–287. 10.1016/j.jtherbio.2011.04.001

Jin, J., Armstrong, R., & Tang, C. (2019). Impact of elevated CO2 on grain nutrient concentration varies with crops and soils – A long-term FACE study. Science of the Total Environment, 651, 2641–2647. 10.1016/j.scitotenv.2018.10.170

Kellermann, V., & van Heerwaarden, B. (2019). Terrestrial insects and climate change: adaptive responses in key traits. In Physiological Entomology (Vol. 44, Issue 2). 10.1111/phen.12282

Kingsolver, J. G., & Huey, R. B. (2008). Size, temperature, and fitness: Three rules. Evolutionary Ecology Research, 10(2), 251–268.

Kreuzwieser, J., & Gessler, A. (2010). Global climate change and tree nutrition: Influence of water availability. Tree Physiology, 30(9), 1221–1234. 10.1093/treephys/tpq055

Kristensen, T. N., Overgaard, J., Lassen, J., Hoffmann, A. A., & Sgrò, C. (2015). Low evolutionary potential for egg-to-adult viability in Drosophila melanogaster at high temperatures. Evolution, 69(3), 803–814. 10.1111/evo.12617

Kutz, T. C., Sgrò, C. M., & Mirth, C. K. (2019). Interacting with change: Diet mediates how larvae respond to their thermal environment. Functional Ecology, 33(10), 1940–1951. 10.1111/1365-2435.13414

Labarbera, M. (1989). Analyzing body size as a factor in ecology and evolution. Annual Review of Ecology and Systematics. *Vol.* 20, 22, 97–117. 10.1146/annurev.ecolsys.20.1.97

Lafuente, E., Duneau, D., & Beldade, P. (2018). Genetic basis of thermal plasticity variation in Drosophila melanogaster body size. PLoS Genetics, 14(9), 1–24. 10.1371/journal.pgen.1007686

Lande, R. (1980). Sexual dimorphism, sexual selection, and adaptation in polygenic characters. Evolution, 34(2), 292. 10.2307/2407393

Lande, R. (2009). Adaptation to an extraordinary environment by evolution of phenotypic plasticity and genetic assimilation. Journal of Evolutionary Biology, 22(7), 1435–1446. 10.1111/j.1420-9101.2009.01754.x

Lasne, C., Hangartner, S. B., Connallon, T., & Sgrò, C. M. (2018). Cross-sex genetic correlations and the evolution of sex-specific local adaptation: Insights from classical trait clines in Drosophila melanogaster. Evolution, 72(6), 1317–1327. 10.1111/evo.13494

Lee, K. P., Kim, J. S., & Min, K. J. (2013). Sexual dimorphism in nutrient intake and life span is mediated by mating in Drosophila melanogaster. Animal Behaviour, 86(5), 987–992. 10.1016/j.anbehav.2013.08.018

Lee, K. P., & Roh, C. (2010). Temperature-by-nutrient interactions affecting growth rate in an insect ectotherm. Entomologia Experimentalis et Applicata, 136(2), 151–163. 10.1111/j.1570-7458.2010.01018.x

Lukac, M., Calfapietra, C., Lagomarsino, A., & Loreto, F. (2010). Global climate change and tree nutrition: Effects of elevated CO2 and temperature. Tree Physiology, 30(9), 1209– 1220. 10.1093/treephys/tpq040

Mackay, T. F. C. (2010). Mutations and quantitative genetic variation : lessons from Drosophila. 2009, 1229–1239. 10.1098/rstb.2009.0315

MacKay, T. F. C., Richards, S., Stone, E. A., Barbadilla, A., Ayroles, J. F., Zhu, D., Casillas, S., Han, Y., Magwire, M. M., Cridland, J. M., Richardson, M. F., Anholt, R. R. H., Barrón, M., Bess, C., Blankenburg, K. P., Carbone, M. A., Castellano, D., Chaboub, L., Duncan, L., … Gibbs, R. A. (2012). The Drosophila melanogaster Genetic Reference Panel. Nature, 482(7384), 173–178. 10.1038/nature10811

Masson-Delmotte, V., Zhai, P., Pörtner, H.-O., Roberts, D., Skea, J., Shukla, P. R., Pirani, A., Moufouma-Okia, W., Péan, C., Pidcock, R., Connors, S., Matthews, J. B. R., Chen, Y., Zhou, X., Gomis, M. I., Lonnoy, E., Maycock, T., Tignor, M., & Waterfield, T. (2018). Intergovernmental panel on climate change-global warming of 1.5 degrees Celcius. In IPCC-Summary for Policymakers. https://report.ipcc.ch/sr15/pdf/sr15_spm_final.pdf%0Ahttp://www.ipcc.ch/report/sr15/

Matavelli, C., Carvalho, M. J. A., Martins, N. E., & Mirth, C. K. (2015). Differences in larval nutritional requirements and female oviposition preference reflect the order of fruit colonization of Zaprionus indianus and Drosophila simulans. Journal of Insect Physiology, 82, 66–74. 10.1016/j.jinsphys.2015.09.003

Midgley, G., & Hannah, L. (2019). Extinction risk from climate change. Biodiversity and Climate Change: Transforming the Biosphere, 294–296. 10.2307/j.ctv8jnzw1.37

Min, K. J., Flatt, T., Kulaots, I., & Tatar, M. (2007). Counting calories in Drosophila diet restriction. Experimental Gerontology, 42(3), 247–251. 10.1016/j.exger.2006.10.009

Mossman, J. A., Biancani, L. M., Zhu, C. T., & Rand, D. M. (2016). Mitonuclear epistasis for development time and its modification by diet in Drosophila. Genetics, 203(1), 463–484. 10.1534/genetics.116.187286

Noer, N. K., Pagter, M., Bahrndorff, S., Malmendal, A., & Kristensen, T. N. (2020). Impacts of thermal fluctuations on heat tolerance and its metabolomic basis in Arabidopsis thaliana, Drosophila melanogaster, and Orchesella cincta. PLoS ONE, *15*(10 October), 1–20. 10.1371/journal.pone.0237201

Ørsted, M., Rohde, P. D., Hoffmann, A. A., Sørensen, P., & Kristensen, T. N. (2018). Environmental variation partitioned into separate heritable components. Evolution, 72(1), 136–152. 10.1111/evo.13391

Partridge, L., Barrie, B., Fowler, K., & French, V. (1994). Evolution and development of body size and cell size in Drosophila melanogaster in response to temperature. Evolution, 48(4), 1269–1276. 10.1111/j.1558-5646.1994.tb05311.x

Pecl, G. T., Araújo, M. B., Bell, J. D., Blanchard, J., Bonebrake, T. C., Chen, I.-C., Clark, T. D., Colwell, R. K., Danielsen, F., Evengård, B., Falconi, L., Ferrier, S., Frusher, S., Garcia, R. A., Griffis, R. B., Hobday, A. J., Janion-Scheepers, C., Jarzyna, M. A., Jennings, S., … Williams, S. E. (2017). Biodiversity redistribution under climate change: Impacts on ecosystems and human well-being. Science, 355(6332), 1–9. 10.1126/science.aai9214

Poissant, J., Wilson, A. J., & Coltman, D. W. (2010). Sex-specific genetic variance and the evolution of sexual dimorphism: A systematic review of cross-sex genetic correlations. Evolution, 64(1), 97–107. 10.1111/j.1558-5646.2009.00793.x

Polak, M., Kroeger, D. E., Cartwright, I. L., & Ponce DeLeon, C. (2004). Genotype-specific responses of fluctuating asymmetry and of preadult survival to the effects of lead and temperature stress in Drosophila melanogaster. Environmental Pollution, 127(1), 145–155. 10.1016/S0269-7491(03)00238-0

Pottier, P., Burke, S., Drobniak, S. M., Lagisz, M., & Nakagawa, S. (2021). Sexual (in)equality? A meta-analysis of sex differences in thermal acclimation capacity across ectotherms. In Functional Ecology (Vol. 35, Issue 12). 10.1111/1365-2435.13899

Reed, T. E., Robin, S. W., Schindler, D. E., Hard, J. J., & Kinnison, M. T. (2010). Phenotypic plasticity and population viability: The importance of environmental predictability. Proceedings of the Royal Society B: Biological Sciences, 277(1699), 3391–3400. 10.1098/rspb.2010.0771

Reeve, J. P., & Fairbairn, D. J. (1996). Sexual size dimorphism as a correlated response to selection on body size: An empirical test of the quantitative genetic model. Evolution, 50(5), 1927–1938. 10.1111/j.1558-5646.1996.tb03580.x

Rho, M. S., & Lee, K. P. (2017). Temperature-driven plasticity in nutrient use and preference in an ectotherm. Oecologia, 185(3), 401–413. 10.1007/s00442-017-3959-4

Richardson, S. J., Press, M. C., Parsons, A. N., & Hartley, S. E. (2002). How do nutrients and warming impact on plant communities and their insect herbivores? A 9-year study from a sub-Arctic heath. Journal of Ecology, 90(3), 544–556. 10.1046/j.1365-2745.2002.00681.x

Rodrigues, L. R., McDermott, H. A., Villanueva, I., Djukarić, J., Ruf, L. C., Amcoff, M., & Snook, R. R. (2022). Fluctuating heat stress during development exposes reproductive costs and putative benefits. Journal of Animal Ecology, 91(2), 391–403. 10.1111/1365-2656.13636

Rohlf, F. J. (2015). The tps series of software. Hystrix, 26(1), 1–4. 10.4404/hystrix-26.1-11264

Rohlf, F. J., & Slice, D. (1990). Extensions of the procrustes method for the optimal superimposition of landmarks. Systematic Zoology, 39(1), 40–59. 10.2307/2992207

Rosenblatt, A. E., & Schmitz, O. J. (2016). Climate change, nutrition, and bottom-up and top-down food web processes. Trends in Ecology and Evolution, 31(12), 965–975. 10.1016/j.tree.2016.09.009

Scheiner, S. M. (1993). Genetics and evolution of phenotypic plasticity. Annual Review of Ecology and Systematics, 24(Figure 1), 35–68. 10.1146/annurev.es.24.110193.000343

Schilthuizen, M., & Kellermann, V. (2014). Contemporary climate change and terrestrial invertebrates: Evolutionary versus plastic changes. Evolutionary Applications, 7(1), 56–67. 10.1111/eva.12116

Schmidt, P. S., Matzkin, L., Ippolito, M., & Eanes, W. F. (2005). Geographic variation in diapause incidence, life-history traits, and climatic adaptation in Drosophila melanogaster. Evolution, 59(8), 1721–1732. 10.1111/j.0014-3820.2005.tb01821.x

Schou, M. F., Kristensen, T. N., Pedersen, A., Göran Karlsson, B., Loeschcke, V., & Malmendal, A. (2017). Metabolic and functional characterization of effects of developmental temperature in Drosophila melanogaster. American Journal of Physiology - Regulatory Integrative and Comparative Physiology, 312(2), R211–R222. 10.1152/ajpregu.00268.2016

Screen, J. (2014). Climate change and extremes. January.

Seebacher, F., White, C. R., & Franklin, C. E. (2015). Physiological plasticity increases resilience of ectothermic animals to climate change. Nature Climate Change, 5(1), 61–66. 10.1038/nclimate2457

Sgrò, C. M., & Hoffmann, A. A. (2004). Genetic correlations, tradeoffs and environmental variation. Heredity, 93(3), 241–248. 10.1038/sj.hdy.6800532

Sgrò, C. M., Overgaard, J., Kristensen, T. N., Mitchell, K. A., Cockerell, F. E., & Hoffmann, A. A. (2010). A comprehensive assessment of geographic variation in heat tolerance and hardening capacity in populations of Drosophila melanogaster from Eastern Australia. Journal of Evolutionary Biology, 23(11), 2484–2493. 10.1111/j.1420-9101.2010.02110.x

Sgrò, C. M., Terblanche, J. S., & Hoffmann, A. A. (2016). What can plasticity contribute to Insect responses to climate change? Annual Review of Entomology, 61(DECEMBER 2015), 433–451. 10.1146/annurev-ento-010715-023859

Sheets, H. D. (2014). PCAGen8. Integrated Morphometrics Package Suite (IMP) 8.

Shingleton, A. W., Masandika, J. R., Thorsen, L. S., Zhu, Y., & Mirth, C. K. (2017). The sex-specific effects of diet quality versus quantity on morphology in Drosophila melanogaster. Royal Society Open Science, 4(9). 10.1098/rsos.170375

Silva-Soares, N. F., Nogueira-Alves, A., Beldade, P., & Mirth, C. K. (2017). Adaptation to new nutritional environments: Larval performance, foraging decisions, and adult oviposition choices in Drosophila suzukii. BMC Ecology, 17(1), 1–13. 10.1186/s12898-017-0131-2

Sisodia, S., & Singh, B. N. (2010). Resistance to environmental stress in Drosophila ananassae: Latitudinal variation and adaptation among populations. Journal of Evolutionary Biology, 23(9), 1979–1988. 10.1111/j.1420-9101.2010.02061.x

Stillwell, R. C., Blanckenhorn, W. U., Teder, T., Davidowitz, G., & Fox, C. W. (2010). Sex differences in phenotypic plasticity affect variation in sexual size dimorphism in insects: From physiology to evolution. Annual Review of Entomology, 55(November), 227–245. 10.1146/annurev-ento-112408-085500

Stillwell, R. C., Wallin, W. G., Hitchcock, L. J., & Fox, C. W. (2007). Phenotypic plasticity in a complex world: Interactive effects of food and temperature on fitness components of a seed beetle. Oecologia, 153(2), 309–321. 10.1007/s00442-007-0748-5

Teuschl, Y., Reim, C., & Blanckenhorn, W. U. (2007). Correlated responses to artificial body size selection in growth, development, phenotypic plasticity and juvenile viability in yellow dung flies. Journal of Evolutionary Biology, 20(1), 87–103. 10.1111/j.1420-9101.2006.01225.x

Tun-Lin, W., Burkot, T. R., & Kay, B. H. (2000). Effects of temperature and larval diet on development rates and survival of the dengue vector Aedes aegypti in north Queensland, Australia. Medical and Veterinary Entomology, 14(1), 31–37. 10.1046/j.1365-2915.2000.00207.x

Urban, M. C., Richardson, J. L., & Freidenfelds, N. A. (2014). Plasticity and genetic adaptation mediate amphibian and reptile responses to climate change. Evolutionary Applications, 7(1), 88–103. 10.1111/eva.12114

Van Der Putten, W. H., Macel, M., & Visser, M. E. (2010). Predicting species distribution and abundance responses to climate change: Why it is essential to include biotic interactions across trophic levels. Philosophical Transactions of the Royal Society B: Biological Sciences, 365(1549), 2025–2034. 10.1098/rstb.2010.0037

van Heerwaarden, B., Kellermann, V., & Sgrò, C. M. (2016). Limited scope for plasticity to increase upper thermal limits. Functional Ecology, 30(12), 1947–1956. 10.1111/1365-2435.12687

Via, S., & Conner, J. (1995). Evolution in heterogeneous environments: Genetic variability within and across different grains in Tribouum castaneum. Heredity, 74(1), 80–90. 10.1038/hdy.1995.10

Via, S., Gomulkiewicz, R., De Jong, G., Scheiner, S. M., Schlichting, C. D., & Van Tienderen, P. H. (1995). Adaptive phenotypic plasticity: consensus and controversy. Trends in Ecology & Evolution, 10(5), 212–217. 10.1016/S0169-5347(00)89061-8

Via, S., & Lande, R. (1985). Genotype-environment interaction and the evolution of phenotypic plasticity. Evolution, 39(3), 505–522. 10.1111/j.1558-5646.1985.tb00391.x

Via, S., & Lande, R. (1987). Evolution of genetic variability in a spatially heterogeneous environment: Effects of genotype–environment interaction. Genetical Research, 49(2), 147–156. 10.1017/S001667230002694X

Walter, G. M., Clark, J., Terranova, D., Cozzolino, S., Cristaudo, A., Hiscock, S. J., & Bridle, J. (2023). Hidden genetic variation in plasticity provides the potential for rapid adaptation to novel environments. New Phytologist, 239(1), 374–387. 10.1111/nph.18744

Walters, R. J., Blanckenhorn, W. U., & Berger, D. (2012). Forecasting extinction risk of ectotherms under climate warming: An evolutionary perspective. Functional Ecology, 26(6), 1324–1338. 10.1111/j.1365-2435.2012.02045.x

Wang, J. B., Lu, H. L., & St. Leger, R. J. (2017). The genetic basis for variation in resistance to infection in the Drosophila melanogaster genetic reference panel. PLoS Pathogens, 13(3), 1–33. 10.1371/journal.ppat.1006260

Wang, S., Tan, X. L., Guo, X. J., & Zhang, F. (2013). Effect of temperature and photoperiod on the development, reproduction, and predation of the predatory ladybird Cheilomenes sexmaculata (Coleoptera: Coccinellidae). Journal of Economic Entomology, 106(6), 2621–2629. 10.1603/EC13095

Wiens, J. J. (2016). Climate-related local extinctions are already widespread among plant and animal species. PLoS Biology, 14(12), 1–18. 10.1371/journal.pbio.2001104

Wood, C. W., & Brodie, E. D. (2015). Environmental effects on the structure of the G-matrix. Evolution, 69(11), 2927–2940. 10.1111/evo.12795

