## Supplementary material for "Within population plastic responses to combined thermal-nutritional stress differ from those in response to single stressors, and are genetically independent across traits in both males and females": Fig. S1

**Supplementary materials**

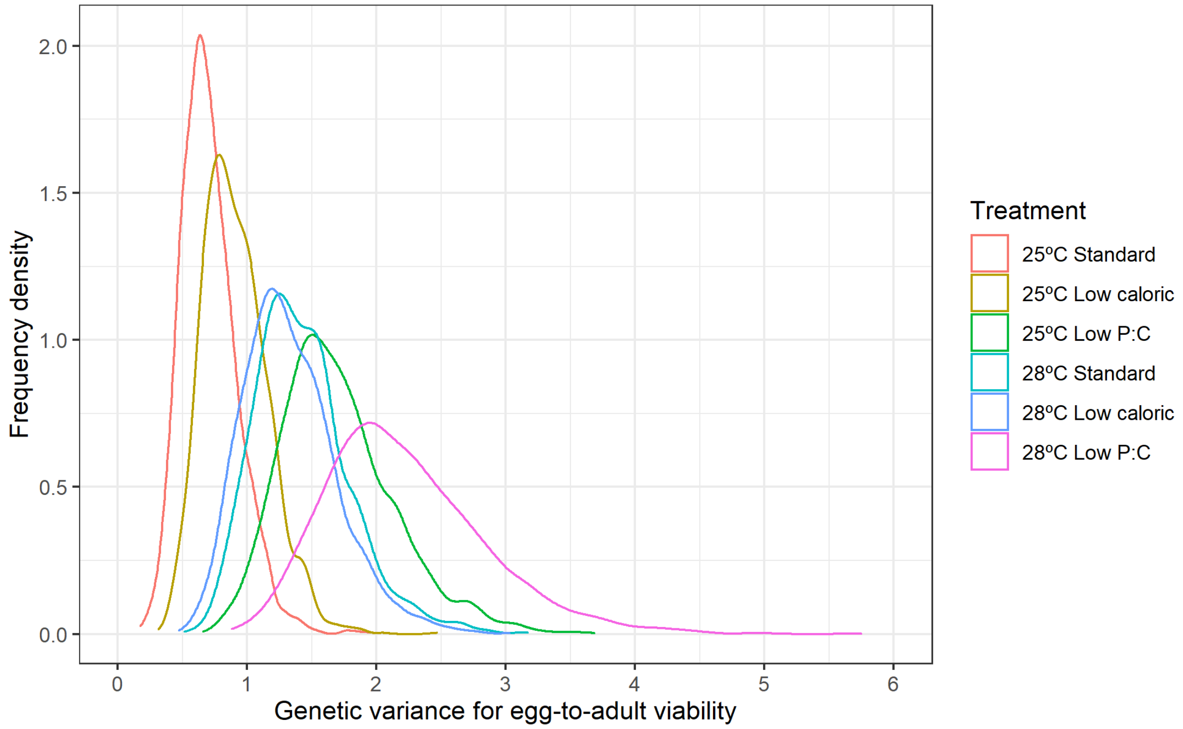


**Fig. S1.** Frequency distributions for the estimates of genetic variance for egg-to-adult viability within each of the six combined thermal-nutritional treatments.

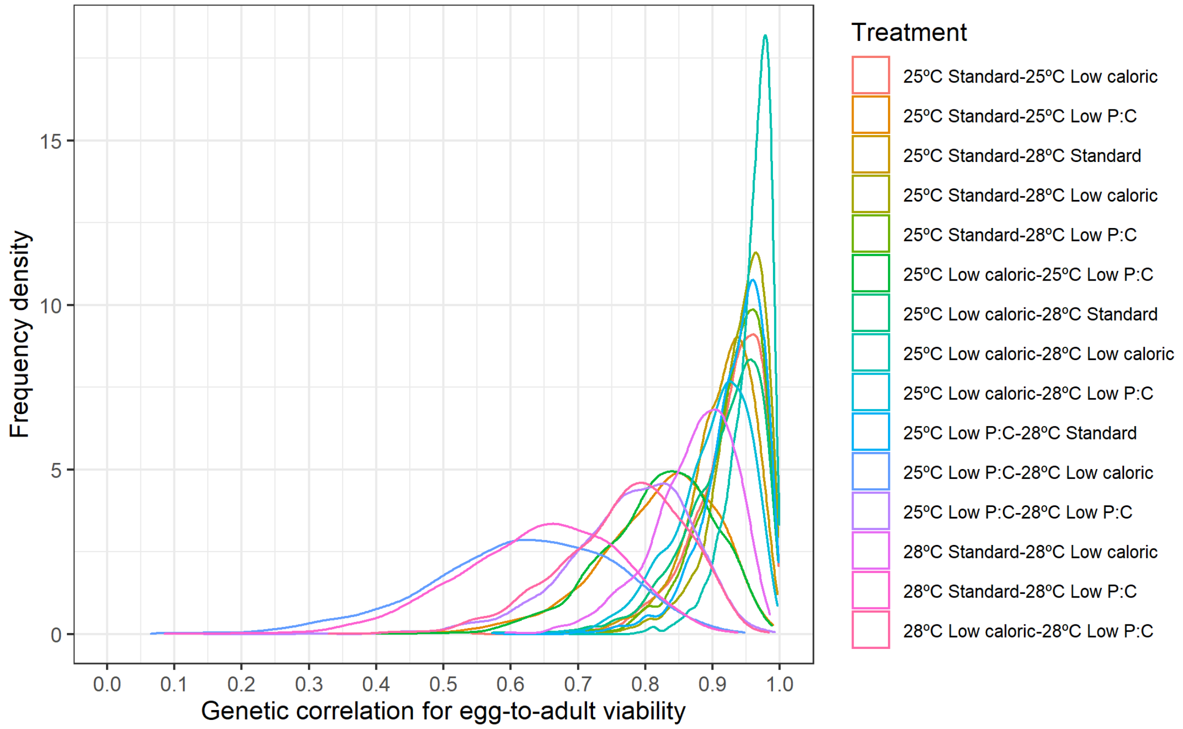


**Fig. S2.** Frequency distributions for the estimates of the cross-environment genetic correlation for egg-to-adult viability between six thermal-nutritional treatments.

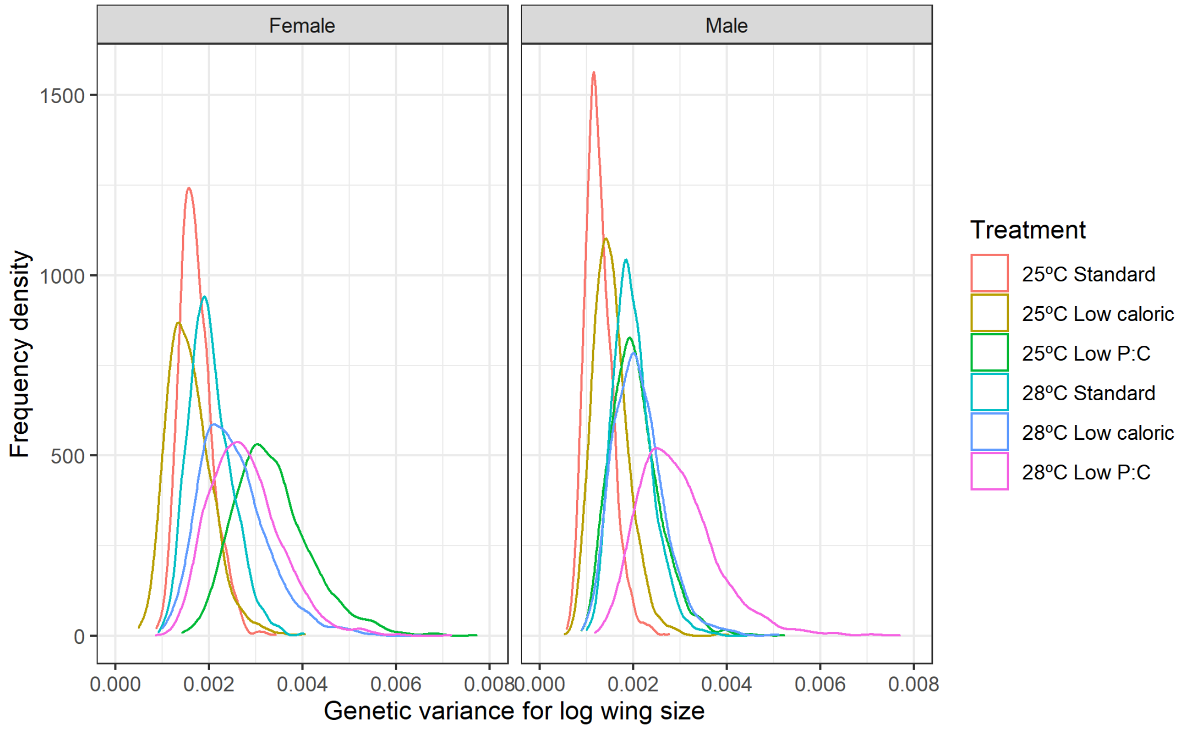


**Fig. S3.** Frequency distributions for genetic variance for log wing size of both sexes within each of the six combined thermal-nutritional treatments.

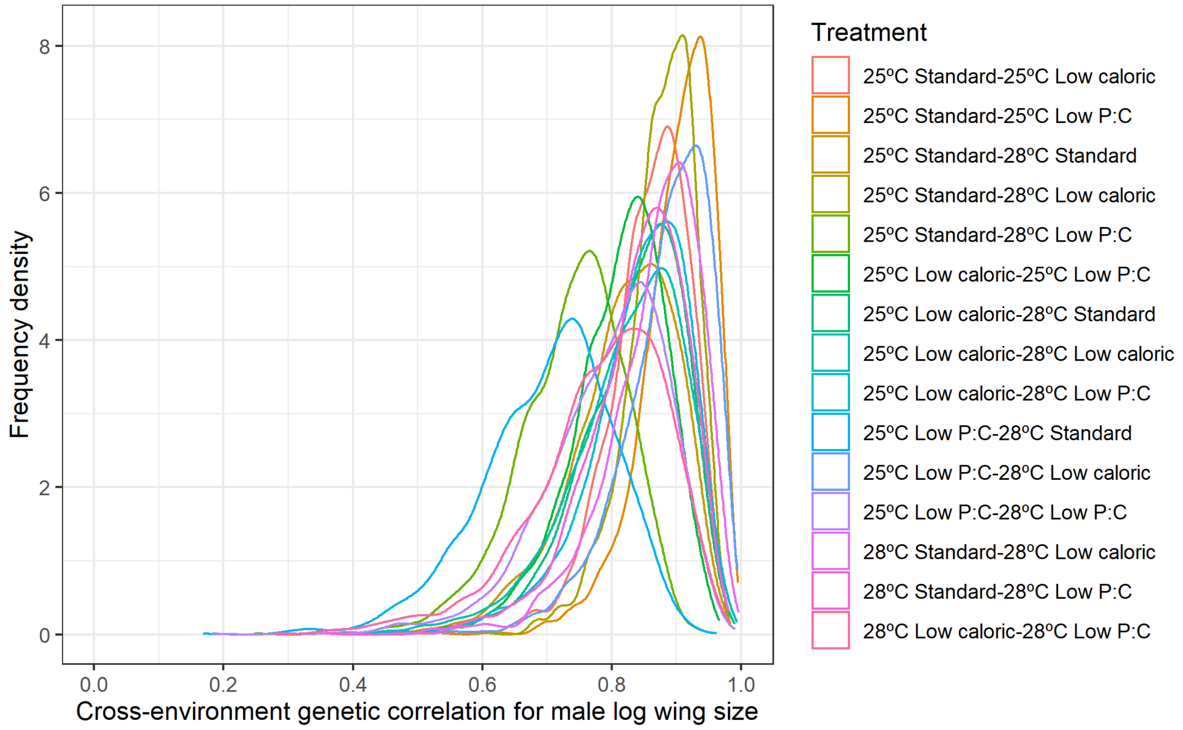


**Fig. S4.** Frequency distributions for estimates of the cross-environment genetic correlation for male log wing size between six thermal-nutritional treatments.

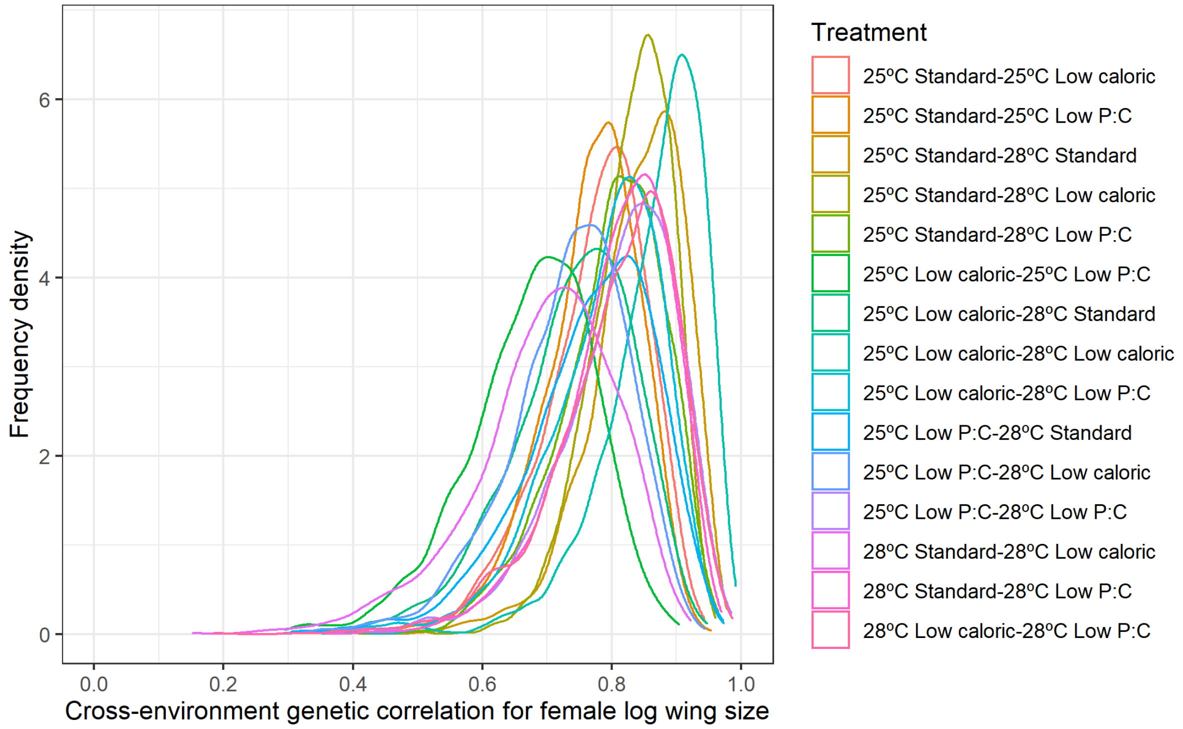


**Fig. S5.** Frequency distributions for the estimates of the cross-environment genetic correlation for female log wing size between six thermal-nutritional treatments.

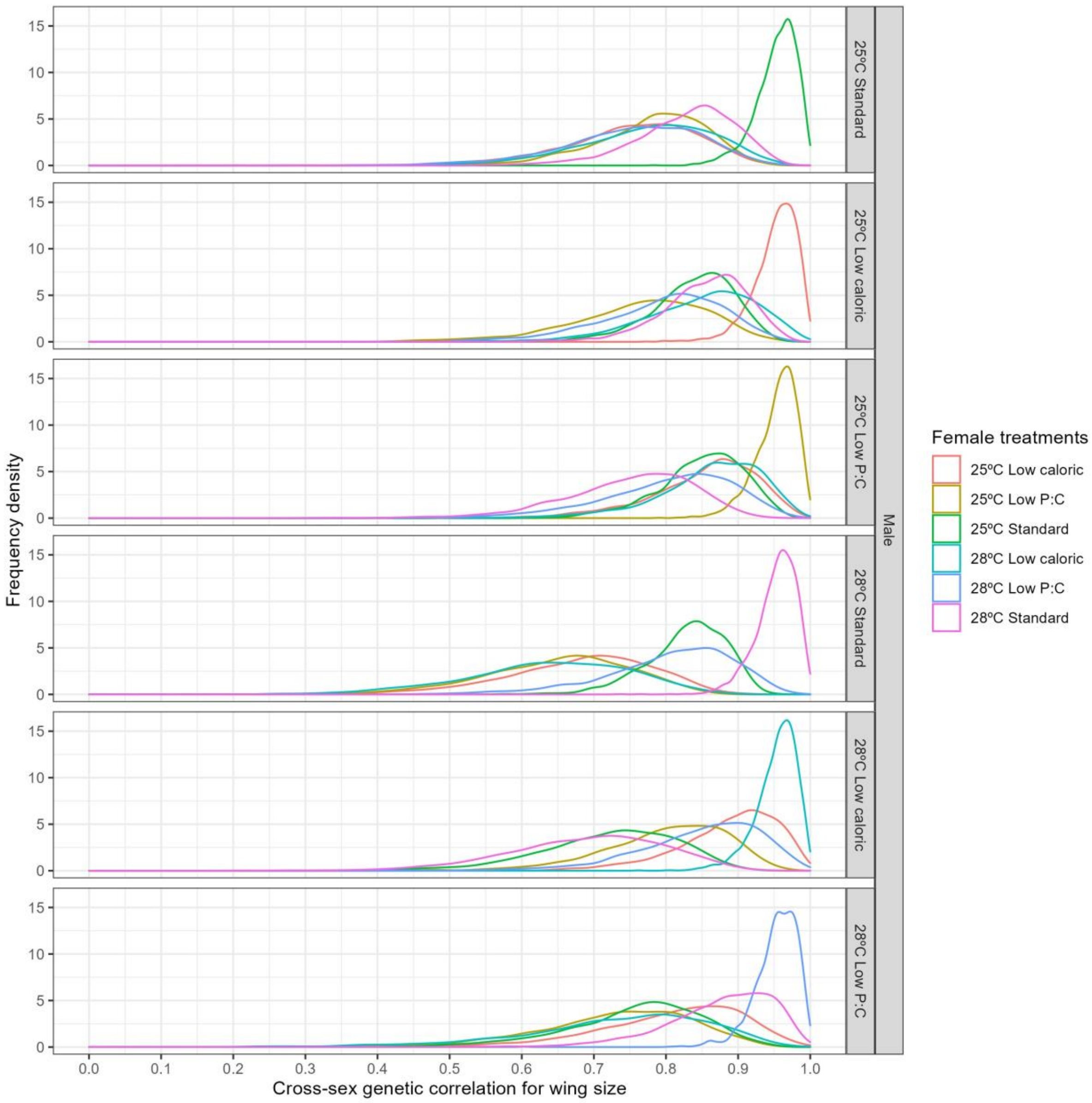


**Fig. S6.** Frequency distributions for the estimates of the cross-sex genetic correlation for wing size between and within six thermal-nutritional treatments.

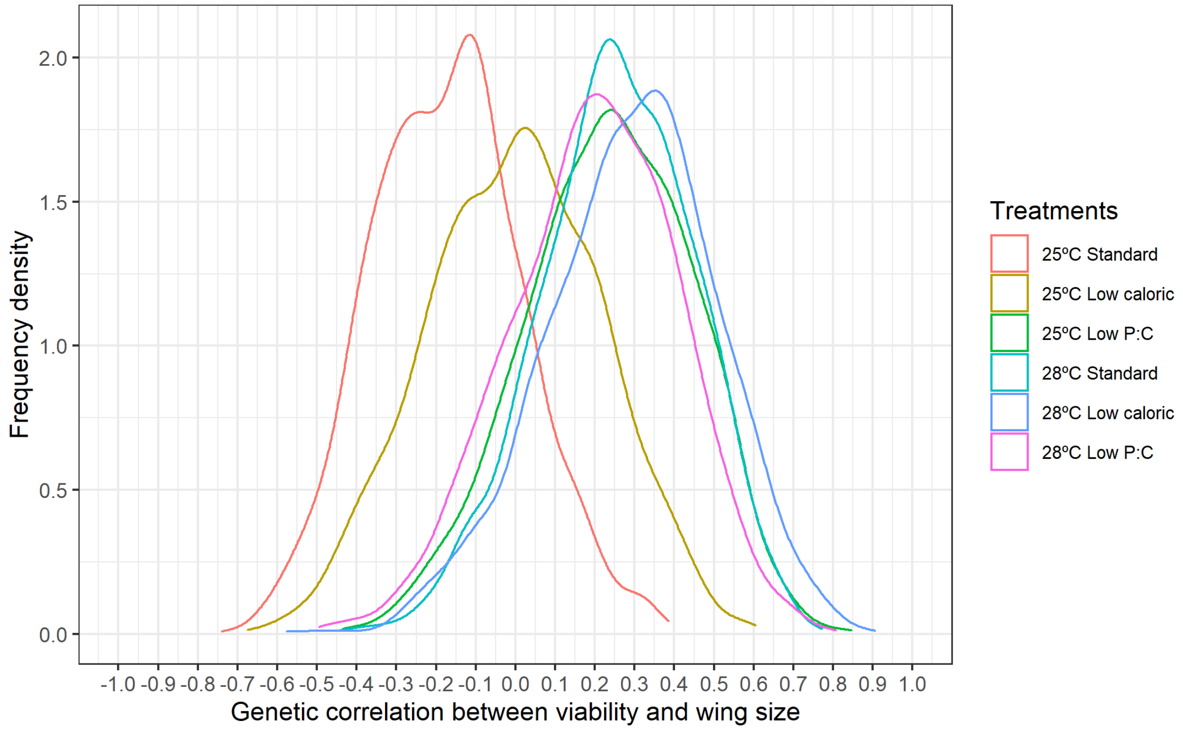


**Fig. S7.** Frequency distributions for the estimates of the genetic correlation between viability and wing size within each of the six experimental treatments.

**Table S1.** The additive genetic covariance matrix for egg-to-adult viability (diagonal: genetic variances in egg-to-adult viability; above the diagonal: genetic correlations among treatments; below the diagonal: genetic covariance among treatments). Numbers in parentheses denote 95% HPD intervals.

| | **25°C Standard** | **25°C Low caloric** | **25°C Low P:C** | **28°C Standard** | **28°C Low caloric** | **28°C Low P:C** |
|---|---|---|---|---|---|---|
| **25°C Standard** | 0.708 (0.357-1.169) | 0.921 (0.815-0.998) | 0.818 (0.643-0.97) | 0.937 (0.851-0.996) | 0.914 (0.802-0.997) | 0.617 (0.328-0.856) |
| **25°C Low caloric** | 0.73 (0.387-1.094) | 0.909 (0.48-1.433) | 0.913 (0.817-0.994) | 0.926 (0.829-0.995) | 0.956 (0.893-0.999) | 0.775 (0.591-0.934) |
| **25°C Low P:C** | 0.889 (0.473-1.388) | 1.126 (0.645-1.662) | 1.703 (0.949-2.677) | 0.819 (0.655-0.967) | 0.898 (0.786-0.991) | 0.872 (0.745-0.975) |
| **28°C Standard** | 0.931 (0.553-1.425) | 1.045 (0.601-1.548) | 1.264 (0.719-1.855) | 1.425 (0.756-2.154) | 0.931 (0.842-0.996) | 0.641 (0.411-0.853) |
| **28°C Low caloric** | 0.876 (0.467-1.331) | 1.042 (0.574-1.527) | 1.339 (0.776-1.96) | 1.271 (0.736-1.849) | 1.329 (0.653-2.012) | 0.768 (0.581-0.927) |
| **28°C Low P:C** | 0.761 (0.33-1.311) | 1.091 (0.54-1.657) | 1.682 (0.934-2.522) | 1.129 (0.525-1.783) | 1.307 (0.691-2.005) | 2.221 (1.139-3.435) |

**Table S2.** The additive genetic covariance matrix for male wing size (diagonal: genetic variances in male wing size; above the diagonal: genetic correlations among treatments; below the diagonal: genetic covariance among treatments). Numbers in parentheses denote 95% HPD intervals.

| | **25°C Standard** | **25°C Low caloric** | **25°C Low P:C** | **28°C Standard** | **28°C Low caloric** | **28°C Low P:C** |
|---|---|---|---|---|---|---|
| **25°C Standard** | 0.00127 (0.000744-0.001848) | 0.859044 (0.741885-0.965064) | 0.901451 (0.788235-0.985617) | 0.875871 (0.766924-0.961552) | 0.832762 (0.671785-0.956848) | 0.884048 (0.742911-0.984404) |
| **25°C Low caloric** | 0.001192 (0.000701-0.001745) | 0.00153 (0.000826-0.002269) | 0.819232 (0.648111-0.956257) | 0.739054 (0.574862-0.886209) | 0.823444 (0.653317-0.974458) | 0.793907 (0.612764-0.951969) |
| **25°C Low P:C** | 0.001448 (0.000842-0.002083) | 0.001442 (0.000809-0.002136) | 0.002054 (0.001137-0.003072) | 0.811963 (0.667807-0.943036) | 0.840998 (0.684793-0.975076) | 0.864843 (0.71361-0.978383) |
| **28°C Standard** | 0.001391 (0.000853-0.001979) | 0.001288 (7e-04-0.001865) | 0.001637 (0.000994-0.002427) | 0.001996 (0.001262-0.002836) | 0.700721 (0.497725-0.879968) | 0.838018 (0.691317-0.971038) |
| **28°C Low caloric** | 0.001357 (0.000782-0.001996) | 0.001471 (0.000789-0.002184) | 0.00174 (0.000997-0.00261) | 0.001433 (0.000771-0.00217) | 0.002117 (0.001152-0.003168) | 0.782189 (0.568626-0.955328) |
| **28°C Low P:C** | 0.001699 (0.001011-0.002563) | 0.001671 (0.000887-0.002543) | 0.002111 (0.001145-0.003158) | 0.002021 (0.001131-0.002938) | 0.001933 (0.001077-0.002976) | 0.002953 (0.001471-0.004598) |

**Table S3.** The additive genetic covariance matrix for female wing size (diagonal: genetic variances in female wing size; above the diagonal: genetic correlations among treatments; below the diagonal: genetic covariance among treatments). Numbers in parentheses denote 95% HPD intervals.

| | **25°C Standard** | **25°C Low caloric** | **25°C Low P:C** | **28°C Standard** | **28°C Low caloric** | **28°C Low P:C** |
|---|---|---|---|---|---|---|
| **25°C Standard** | 0.001724 (0.001097-0.002431) | 0.773978 (0.600858-0.905373) | 0.768436 (0.61841-0.901187) | 0.830309 (0.711281-0.938566) | 0.739975 (0.548687-0.904866) | 0.732779 (0.544262-0.892409) |
| **25°C Low caloric** | 0.001273 (0.000659-0.001909) | 0.001591 (0.000762-0.002588) | 0.840448 (0.702861-0.959872) | 0.800011 (0.637864-0.943489) | 0.868882 (0.722222-0.982045) | 0.810407 (0.636057-0.968463) |
| **25°C Low P:C** | 0.001845 (0.00108-0.002769) | 0.00193 (0.001028-0.002988) | 0.003355 (0.00183-0.00499) | 0.675903 (0.462352-0.837153) | 0.790525 (0.606499-0.931111) | 0.696671 (0.476072-0.898116) |
| **28°C Standard** | 0.001553 (0.000911-0.002221) | 0.00143 (0.000745-0.002149) | 0.001764 (0.000829-0.002714) | 0.002037 (0.00123-0.00296) | 0.767052 (0.556713-0.929885) | 0.808932 (0.638368-0.947558) |
| **28°C Low caloric** | 0.001536 (0.000788-0.002324) | 0.001723 (0.000917-0.002701) | 0.00229 (0.001104-0.00347) | 0.00173 (0.000819-0.002623) | 0.00253 (0.001237-0.004008) | 0.807181 (0.615288-0.96234) |
| **28°C Low P:C** | 0.001599 (0.000861-0.002415) | 0.00169 (0.000753-0.002549) | 0.002129 (0.001011-0.003445) | 0.001919 (0.001065-0.002905) | 0.002126 (0.001042-0.003276) | 0.002784 (0.001405-0.004188) |

**Table S4.** Cross-sex genetic correlation of wing size between and within treatments. Numbers in parentheses denote 95% HPD intervals.

| Female / Male | 25°C Standard | 25°C Low caloric | 25°C Low P:C | 28°C Standard | 28°C Low caloric | 28°C Low P:C |
|---|---|---|---|---|---|---|
| **25°C Standard** | 0.953089 (0.896584-0.997699) | 0.751245 (0.564683-0.918565) | 0.779705 (0.630725-0.91648) | 0.827786 (0.688606-0.947673) | 0.775122 (0.590367-0.941071) | 0.751162 (0.551401-0.923017) |
| **25°C Low caloric** | 0.837796 (0.728236-0.948437) | 0.951987 (0.894851-0.997544) | 0.763301 (0.564538-0.931164) | 0.853992 (0.738263-0.957606) | 0.847465 (0.694024-0.979265) | 0.793931 (0.618203-0.94087) |
| **25°C Low P:C** | 0.849296 (0.72922-0.955584) | 0.852147 (0.692403-0.971712) | 0.952873 (0.897657-0.998584) | 0.750338 (0.562473-0.892509) | 0.861491 (0.714681-0.982559) | 0.802273 (0.615639-0.955194) |
| **28°C Standard** | 0.829143 (0.719843-0.921201) | 0.680949 (0.479185-0.863783) | 0.650169 (0.454854-0.838144) | 0.952561 (0.898986-0.998201) | 0.636473 (0.405641-0.836235) | 0.805617 (0.628011-0.95811) |
| **28°C Low caloric** | 0.72214 (0.539914-0.891487) | 0.88539 (0.745469-0.994507) | 0.800184 (0.635798-0.950226) | 0.690932 (0.490708-0.89253) | 0.952911 (0.897236-0.997323) | 0.847993 (0.684783-0.988736) |
| **28°C Low P:C** | 0.758806 (0.565261-0.923945) | 0.807421 (0.611-0.976932) | 0.728527 (0.509774-0.913564) | 0.875463 (0.730603-0.987041) | 0.733701 (0.475757-0.949508) | 0.95316 (0.899002-0.997566) |

**Table S5.** Genetic correlation between egg-to-adult viability and wing size in each treatment.

| Treatment | Correlation | Lower HPD | Upper HPD |
|---|---|---|---|
| 25°C Standard | -0.173 | -0.554 | 0.186 |
| 25°C Low caloric | -0.006 | -0.431 | 0.417 |
| 25°C Low P:C | 0.234 | -0.191 | 0.612 |
| 28°C Standard | 0.253 | -0.127 | 0.617 |
| 28°C Low caloric | 0.29 | -0.169 | 0.675 |
| 28°C Low P:C | 0.195 | -0.193 | 0.62 |
